# Consciousness is supported by near-critical cortical electrodynamics

**DOI:** 10.1101/2021.06.10.447959

**Authors:** Daniel Toker, Ioannis Pappas, Janna D. Lendner, Joel Frohlich, Diego M. Mateos, Suresh Muthukumaraswamy, Robin Carhart-Harris, Michelle Paff, Paul M. Vespa, Martin M. Monti, Friedrich T. Sommer, Robert T. Knight, Mark D’Esposito

## Abstract

Mounting evidence suggests that during conscious states, the electrodynamics of the cortex are poised near a critical point or phase transition, and that this near-critical behavior supports the vast flow of information through cortical networks during conscious states. Here, for the first time, we empirically identify the specific critical point near which conscious cortical dynamics operate as the edge-of-chaos critical point, or the boundary between periodicity/stability and chaos/instability. We do so by applying the recently developed modified 0-1 chaos test to electrocorticography (ECoG) and magnetoencephalography (MEG) recordings from the cortices of humans and macaques across normal waking, generalized seizure, GABAergic anesthesia, and psychedelic states. Our evidence suggests that cortical information processing is disrupted during unconscious states because of a transition of cortical dynamics away from this critical point; conversely, we show that psychedelics may increase the information-richness of cortical activity by tuning cortical electrodynamics closer to this critical point. Finally, we analyze clinical electroencephalography (EEG) recordings from patients with disorders of consciousness (DOC), and show that assessing the proximity of cortical electrodynamics to the edge-of-chaos critical point may be clinically useful as a new biomarker of consciousness.

**Significance Statement:** What changes in the brain when we lose consciousness? One possibility is that the loss of consciousness corresponds to a transition of the brain’s electric activity away from edge-of-chaos criticality, or the knife’s edge in between stability and chaos. Recent mathematical developments have produced novel tools for testing this hypothesis, which we apply for the first time to cortical recordings from diverse brain states. We show that the electric activity of the cortex is indeed poised near the boundary between stability and chaos during conscious states and transitions away from this boundary during unconsciousness, and that this transition disrupts cortical information processing.

## Introduction

What are the dynamical properties of electric brain activity that are necessary for consciousness, and how are those properties disrupted during unconscious states such as surgical anesthesia, generalized seizures, coma, and vegetative states?

One possibility, which is suggested by a large body of recent evidence, is that the electrodynamics of the conscious brain are poised near some sort of phase transition or “critical point,” and that this near-critical behavior supports the vast flow o f information through the brain during conscious states (1, 2). A critical point refers to the knife’s edge in between different phases of a system (e.g. liquid to solid water) or types of dynamical states (e.g. laminar to turbulent airflow). It is widely believed that electrodynamics of both micro- and macro-scale cortical networks are poised near some critical point, because power-law statistics, which are a key signature of criticality (3), are consistently identified in recordings of neural electrodynamics (4, 5). And such critical behavior is known to have important computational benefits: because critical and near-critical systems tend to have a high capacity for encoding and transmitting information (6–9), it is widely believed that being poised at - or at least *near* (10, 11) - criticality of some form endows neural populations with a high capacity for encoding sensory signals and for communicating with other neural populations (4, 5, 12), particularly during conscious states (1, 2). On the flip side, because signatures of cortical criticality have been observed to disappear or diminish during unconscious states (4, 13, 14), it may be that a transition of cortical activity away from criticality is what underlies the disruption to cortical information processing during unconscious states (2).

Though the existing evidence supports this conjectured relationship between criticality, cortical information processing, and conscious vs. unconscious brain states, prior empirical work has, for the most part, relied on the detection of power-law statistics in neural electrodynamics, most typically in the form of “neuronal avalanches” or bursts of electric activity whose sizes follow a power-law distribution, in order to infer neural criticality during conscious states and a loss of criticality during unconscious states (15); but, the detection of power-law statistics alone cannot specify the type of critical point a system is poised at, because power-law statistics appear across many types of phase transitions (3). Moreover, neuronal avalanches can arise in non-critical neural systems (16), and neural networks can display several unique dynamical critical points, only one of which is the phase transition that gives rise to neuronal avalanches (17). Though some prior studies have attempted to use alternative metrics to assess the relationship between neural criticality and consciousness (18–20), the precise form of criticality under consideration has largely remained mathematically unspecified (15), which leaves open the fundamental question: what, exactly, is this phase transition near which cortical electrodynamics seem to operate during conscious states? Put another way: what, from a mathematical perspective, are the dynamical phases that lie on either side of this critical point? Terms like “order” and “disorder” have commonly been used to describe the phases on either side of neural criticality, but these terms are imprecise unless they are defined relative to the breaking of a specific form of mathematical symmetry, where the “ordered” phase of a system is the symmetry-broken phase (in the way that ice is the "ordered" phase of water relative to the freezing critical point, because water loses its translational and rotational symmetry at this phase transition) - see SI Appendix, Supplementary Note 1 for a more detailed discussion of this point. Imprecise use of terms like “order” and “disorder” can also be misleading in the context of neural criticality. For example, chaos, which is defined as exponential sensitivity to small perturbations, is often used interchangeably with “disorder” in the literature on neural criticality (15), but chaos is in fact the “ordered” phase of dynamical systems because it corresponds to the breaking of topological or de-Rahm supersymmetry (21) (SI Appendix, Supplementary Note 1). This inconsistency and lack of mathematical specificity in definitions of neural criticality may underlie the apparent variability of prior results relating criticality to different brain states, where, for example, some purported metrics of criticality seem to suggest that seizures constitute a departure from criticality while others seem to suggest that seizures are in fact critical phenomena (22). If, as has been proposed (1), the disruption to cortical information processing during unconscious states is mediated by an excursion of cortical dynamics away from some sort of critical point during these states, then mathematically precise identification of this critical point may be crucial for improving both our theoretical and clinical grasp on the neural correlates of consciousness.

Here, we provide the first direct empirical evidence for the hypothesis (23) that during conscious states, cortical electrodynamics specifically operate near a mathematically well-defined critical point known as edge-of-chaos criticality, which is the phase transition from periodic/stable to chaotic/unstable dynamics. Many systems (6–9, 24), including deep neural networks (24) and echo state networks (8), have been shown to exhibit their highest capacity for information processing precisely at this specific critical point. In line with this well-replicated phenomenon, we show that excursions of low-frequency cortical activity away from this critical point during generalized seizures and GABAergic anesthesia induce both a loss of information in cortical dynamics as well as a loss of consciousness. We moreover show that lysergic acid diethylamide (LSD), a 5-HT_2*A*_ receptor agonist characterized as a hallucinogen or “psychedelic,” may tune cortical dynamics closer to the edge-of-chaos critical point relative to normal waking states, which increases the information-richness of cortical activity. Finally, we provide preliminary evidence that cortical electrodynamics return to the vicinity of this critical point as patients with disorders of consciousness (DOC) regain awareness, which suggests that assessing the proximity of cortical dynamics to edge-of-chaos criticality may be useful as a new clinical biomarker of consciousness. We provide Matlab (R2020a) code for our analysis in the hopes of facilitating further basic and translational research along these lines.

## Results

### Mean-field dynamics

To empirically assess whether cortical dynamics operate near the edge-of-chaos critical point during conscious states, and whether this underpins the information-richness of cortical dynamics during conscious states (Fig. 1), we must first assess varying levels of chaoticity and information-richness in a model of cortical electrodynamics, and then test whether real data agree with the model’s predictions. The reason we must first analyze a model is because a system’s level of stability can only be detected with certainty in a simulation, where noise and initial conditions can be precisely controlled. For this reason, it is generally agreed (25) that empirical evidence of varying levels of chaos in a biological system requires comparison of real data to an accurate model of the biological system of interest. Toward that end, we assessed the mean-field model of macro-scale cortical electrodynamics developed by Steyn-Ross, Steyn-Ross, and Sleigh (26) because it has been shown to successfully model the low-frequency macro-scale cortical electrodynamics of waking conscious (26), generalized seizure (26–28), and GABAergic anesthesia (26, 29) states, and thus can be compared to real recordings of large-scale cortical electrodynamics across these diverse brain states. The model is also unique in its inclusion of gap junction coupling between cortical interneurons, which recent empirical work in zebrafish has shown is an important mechanism for the maintenance of criticality in electric neural activity (30). Using this model, we generated 10-second simulations of macro-scale cortical electrodynamics corresponding to waking conscious, generalized seizure, and GABAergic anesthesia states (using parameter ranges identified in past studies - see Materials and Methods), and additionally performed a parameter sweep on the model to generate dynamics from 773 non-biologically specific states in order to more broadly assess the relationship between proximity to edge-of-chaos criticality and information-richness (see Materials and Methods). For each biologically specific and n on-biologically specific state, we calculated the largest Lyapunov exponent, which is a mathematically formal measure of chaoticity that can only be accurately estimated in simulations, of the deterministic component of the model’s dynamics (i.e., with the model’s noise inputs turned off - see Materials and Methods). Note that a largest Lyapunov exponent of 0 corresponds to edge-of-chaos criticality, a positive largest Lyapunov exponent corresponds to chaos/instability, and a negative largest Lyapunov exponent corresponds to periodicity/stability. Finally, to assess the information-richness of the model’s behavior, we calculated the Lempel-Ziv complexity of its full dynamics (with noise inputs turned on) using three variants of Lempel-Ziv complexity (see Materials and Methods). As a measure of the compressibility of a time-series (31), Lempel-Ziv complexity directly quantifies the amount of non-redundant information in a time-series, as compressibility is mathematically lower-bounded by the amount of unique information in a signal (32). While there are several measures of information-richness (e.g. Shannon entropy), we here use Lempel-Ziv complexity both because it can be accurately estimated from short, noisy, nonlinear time-series, and because the Lempel-Ziv complexity of cortical electrodynamics has been shown to consistently drop during unconscious states - see Frohlich et al (33) for an in-depth discussion of the relationship between Lempel-Ziv complexity and consciousness, including a critical assessment of purported dissociations between Lempel-Ziv complexity and conscious vs. unconscious brain states.

**Fig. 1.**
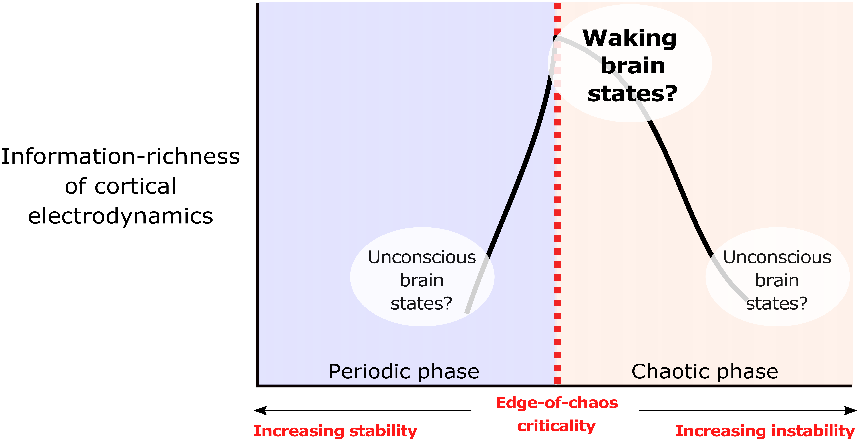
Hypothesized relationship between consciousness, edge-of-chaos criticality, and cortical information processing. We suggest that the electrodynamics of the cortex may be poised near the edge-of-chaos critical point during conscious states, and transition away from this specific critical point during unconscious states. According to this hypothesis, transitions of cortical electrodynamics away from this critical point - either into the chaotic phase (leading to dynamical instability) or into the periodic phase (leading to hyper-stability) - should disrupt cortical information processing and induce unconsciousness. In other words, we should expect to see an inverse-U relationship between chaoticity and information processing in the cortex, with cortical dynamics during conscious states near the top of this inverse-U (i.e., in the near-critical, information-rich regime), and we should moreover expect to see cortical dynamics during unconscious states at either the bottom right of this inverse-U (i.e., the unstable, information-poor regime) or at the bottom left of this inverse-U (i.e., the hyper-stable, information-poor regime) (1, 2, 21). Such an inverse-U relationship between chaoticity and information processing has been observed in many other dynamical systems (6–9), but remains to be empirically observed in the brain.

Consistent with the prediction that the cortex generates information-rich dynamics during conscious states by operating near the edge-of-chaos critical point, we found that the Lempel-Ziv complexity of the model’s simulated electrodynamics (with noise inputs) was maximal when the deterministic component of its dynamics were poised near this critical onset of chaos (red vertical line in Fig. 2A), and that the model’s simulation of the conscious, waking state was near this critical, information-rich regime. The model specifically placed waking, conscious cortical dynamics on the chaotic/unstable side of this critical edge (black circle in Fig. 2A). Moreover, as predicted, the model exhibited a general inverse-U relationship between chaoticity and information-richness, with the amount of non-redundant information generated by its dynamics falling both in the chaotic/unstable phase (bottom right of the inverse-U) and in the periodic/stable phase (bottom left of the inverse-U), similar to what has been shown in many other systems (6–9, 34). To quantitatively confirm this qualitative result, we used Simonsohn’s two lines statistical test of a U-shaped relationship, which accepts a null hypothesis of no U-shaped relationship if either of two opposite-sign regression lines (one for high and one for low values of the *x* variable) are statistically insignificant - see Simonsohn (35) for details on this test. The two lines test failed to reject the null hypothesis no U-shaped relationship between largest Lyapunov exponents and univariate, joint, or concatenated Lempel-Ziv complexity (Table 1). Finally, we note that the mean-field model specifically placed GABAergic anesthesia in the strongly chaotic/unstable phase and placed generalized seizures in the periodic/stable phase, even though both simulated states led to information loss (Fig. 2A) and increased spectral power at low frequencies (SI Appendix, Fig. S1).

**Fig. 2.**
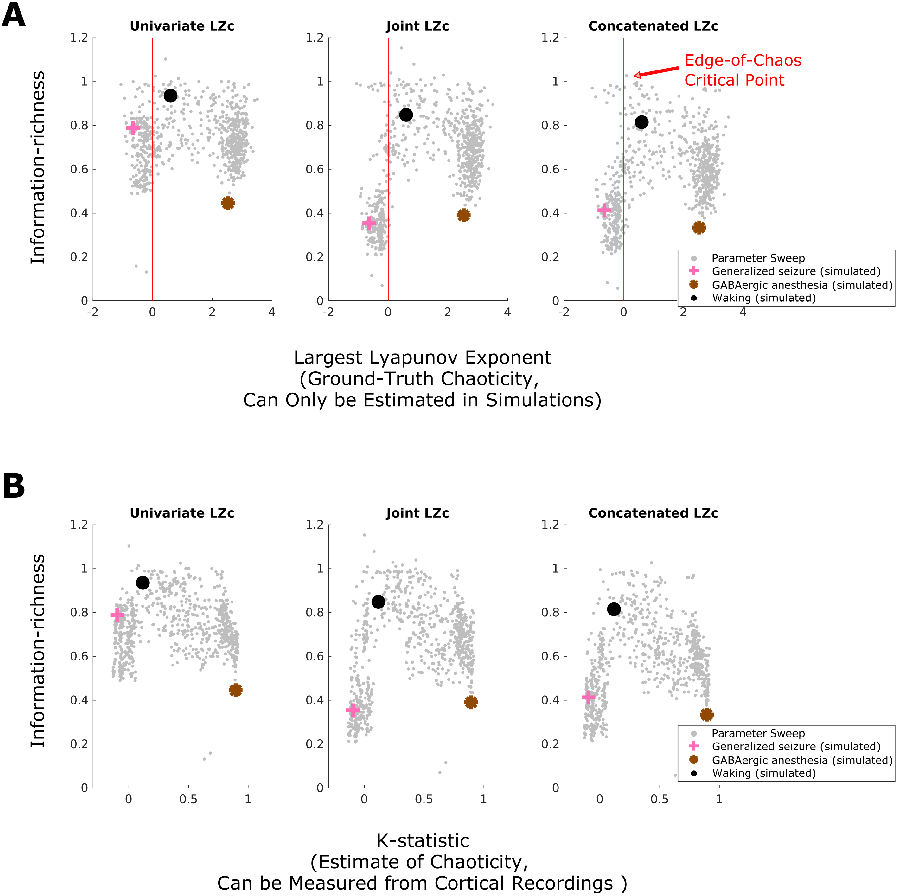
Predictions relating brain states, information processing, and the criticality of low-frequency cortical electrodynamics, and the testability of those predictions in real data. **A** We calculated both the largest Lyapunov Exponent (ground-truth instability) and Lempel-Ziv complexity (information-richness) of 10-second mean-field simulations of low-frequency cortical electrodynamics during waking conscious (black circle), generalized seizure (pink cross), and GABAergic anesthesia (brown asterisk) states. We also performed a parameter sweep of the mean-field model to more generally assess the relationship between the information-richness of its dynamics and the proximity of those dynamics to this critical point (see Materials and Methods); each small gray dot represents the result of a single 10-second simulation with a unique parameter configuration that did not correspond to a biologically specific brain state. We found that all three measures of information-richness peak near the edge-of-chaos critical point (red vertical line), and that the simulated waking conscious dynamics are near this critical, information-rich regime. Importantly, waking cortical dynamics are here predicted to lie on the unstable side of this critical point. All three information measures drop in both the chaotic/unstable phase (positive largest Lyapunov exponent), where GABAergic anesthesia cortical dynamics are predicted to lie, and also in the periodic/stable phase (negative largest Lyapunov exponent), where generalized seizure dynamics are predicted to lie. **B** The modified 0-1 chaos test (see Materials and Methods), when applied to the low-pass filtered simulated dynamics of the mean-field model, accurately tracks the chaoticity of those dynamics and is able to recapitulate the ground-truth inverse-U relationship between chaoticity and information-richness. This validates the ability of the modified 0-1 chaos test to empirically evaluate these specific predictions relating consciousness, information processing, and the proximity of low-frequency cortical electrodynamics to the edge-of-chaos critical point in real cortical recordings.

**Table 1.** Results of Simonsohn’s Two-Lines Test of a U-shaped relationship (35). The test confirmed the U-shaped relationship (across different states of the mean-field model of cortical electrodynamics) between all three measures of Lempel-Ziv complexity (LZc) and chaoticity, as measured by both ground-truth largest Lyapunov exponents (LLE) and the K-statistic of the modified 0-1 chaos test. The test also confirmed the U-shaped relationship (across subjects) in our cortical recordings between chaoticity, as measured by the K-statistic, and both univariate and concatenated Lempel-Ziv complexity. P-values were Bonferroni-corrected for multiple comparisons against the same set of either largest Lyapunov exponents (LLE) or K-statistic values.

| Simonsohn's Two-Lines Test Results |  |  |
| --- | --- | --- |
|  | Regression Line 1 | Regression Line 2 |
| **Simulation data** |  |  |
| LLE vs Univariate LZc | b=0.1, z=5.65, p< 10<sup>-4</sup> | b=-0.06, z=-5.44, p< 10<sup>-4</sup> |
| LLE vs Joint LZc | b=0.4, z=11, p< 10<sup>-4</sup> | b=-0.08, z=-8.39, p< 10<sup>-4</sup> |
| LLE vs Concatenated LZc | b=0.29, z=8.89, p< 10<sup>-4</sup> | b=-0.08, z=-6.98, p< 10<sup>-4</sup> |
| K vs Univariate LZc | b=0.71, z=11.76, p< 10<sup>-4</sup> | b=-0.28, z=-11.99, p< 10<sup>-4</sup> |
| K vs Joint LZc | b=1.79, z=25.4, p< 10<sup>-4</sup> | b=-0.32, z=-11.66, p< 10<sup>-4</sup> |
| K vs Concatenated LZc | b=1.49, z=21.49, p< 10<sup>-4</sup> | b=-0.36, z=-12.62, p< 10<sup>-4</sup> |
| **Empirical data** |  |  |
| K vs Univariate LZc | b=0.26, z=10.74, p< 10<sup>-4</sup> | b=-1.03, z=-8.42, p< 10<sup>-4</sup> |
| K vs Joint LZc | b=0.12, z=1.99, p=0.137 | b=-1.38, z=-6.55, p< 10<sup>-4</sup> |
| K vs Concatenated LZc | b=0.33, z=6.01, p< 10<sup>-4</sup> | b=-1.25, z=-10, p< 10<sup>-4</sup> |

Such predictions of varying degrees of chaoticity in real biological systems have historically been difficult to test, but recent mathematical developments in nonlinear time-series analysis now allow for accurate detection of chaoticity from noisy time-series data. In particular, the modified 0-1 chaos test has emerged as a robust measure of instability from noisy recordings (25, 36–40) (see Materials and Methods). Given a recorded time-series, the 0-1 chaos test outputs a statistic *K*, which estimates the degree of chaoticity of a (predominantly) deterministic signal on a scale from 0 to 1; lower values indicate periodicity/stability and higher values indicate chaos/instability. In order to specifically assess the chaoticity of low-frequency cortical electrodynamics (as simulated in the mean-field model), we low-pass filtered all time-series data in this study before applying the modified 0-1 chaos test. While low-pass filter cutoffs are often selected at canonical frequency bands, recent work has shown that this approach can induce spurious oscillations when no such oscillations are present, and can moreover obfuscate natural but meaningful variance in oscillation frequencies across channels, subjects, and species; for these reasons, to select low-pass filter cutoffs for every channel in every trial, we used the data-driven “Fitting Oscillations and One Over F” or “FOOOF” algorithm, which helps identify real channel-specific oscillations and their respective frequencies based neural power spectra (41). We then applied the modified 0-1 chaos test to these low-pass filtered signals (see Materials and Methods for more details). In addition, we verified that the majority of signals analyzed in this paper were generated by predominantly deterministic processes (SI Appendix, Tables S1-S2), which is an important assumption of the modified 0-1 chaos test. Finally, where applicable, our statistical analyses included these selected low-pass filter frequencies as a covariate, in order to ensure that our results are driven by the stability of low-frequency cortical oscillations, rather than by their frequencies.

Confirming the ability of the modified 0-1 chaos test to detect varying levels of chaoticity from real time-series data, we found that its *K*-statistic, when applied to the model’s simulated dynamics (with noise inputs turned on) after low-pass filtering using the FOOOF algorithm, was strongly correlated with the ground-truth largest Lyapunov exponent of the deterministic component of the mean-field model’s dynamics (which can only be estimated in simulations) (r=0.84, p<10^−4^ Bonferroni-corrected; partial correlation *ρ*=0.82 after controlling for selected low-pass filter frequencies, p<10^−4^ Bonferroni-corrected), and that this correlation was robust to high levels of both white and pink (1/f) measurement noise (Tables S3-S4). The K-statistic of these low-pass filtered signals was likewise correlated with the stochastic Lyapunov exponents of the model (i.e., with Lyapunov exponents calculated for partially stochastic simulations with identical noise inputs) (r=0.83, p<10^−4^; partial correlation *ρ*=0.81 after controlling for selected low-pass filter frequencies, p<10^−4^). Moreover, the *K*-statistic was able to recapitulate the inverse-U relationship between chaoticity and Lempel-Ziv complexity in the model, as shown qualitatively in Fig. 2B. As was the case for the ground-truth largest Lyapunov exponents, Simonsohn’s two lines test quantitatively confirmed the inverse-U relationship between the *K*-statistic and univariate, joint, and concatenated Lempel-Ziv complexity (Table 1). These results indicate that we can use the 0-1 test’s *K*-statistic to empirically test, for the first time, the above-mentioned predictions relating consciousness, information-richness, and cortical instability relative to the edge-of-chaos critical point in real recordings of macro-scale cortical electrodynamics.

### Cortical electrodynamics confirm mean-field predictions

We therefore applied the modified 0 - 1 chaos test to low-frequency activity extracted from surface electrocorticography (ECoG) recordings of the cortical electrodynamics of two macaques and five human epilepsy patients during normal waking states, of two macaques and three human epilepsy patients under GABAergic (propofol, or propofol and sevoflurane) anesthesia, and of two human epilepsy patients experiencing generalized seizures; we further applied this test to magnetoencephalography (MEG) recordings of the cortical electrodynamics of a third human epilepsy patient experiencing a generalized seizure. We also applied the 0-1 chaos test to the low-frequency component of MEG recordings of the cortical electrodynamics of 16 human subjects under the influence of either a saline placebo or LSD, as psychedelics are the only known compounds to reliably increase the information-richness of cortical electrodynamics (1, 2, 42, 43), and are thought to do so by tuning cortical dynamics closer to some critical point (2, 44). Psychedelics therefore allow us to test a specific and counter-intuitive prediction of this chaos-vs-information processing framework: if cortical electrodynamics during normal waking states do indeed lie on the chaotic side of the edge-of-chaos critical point (as the mean-field model predicts), then psychedelics should, counter-intuitively, increase the information-richness of cortical activity by *reducing* the chaoticity of cortical dynamics, as those dynamics approach the edge-of-chaos critical point from the unstable side of the edge (where normal waking dynamics are predicted to lie).

Confirming our predictions, our empirical analysis yielded an inverse-U relationship between chaoticity and information-richness (as measured by three variants of Lempel-Ziv complexity) in our recordings of cortical electrodynamics, with conscious states at the top of this inverse-U, as shown qualitatively in Fig. 3. To confirm this result quantitatively, we applied Simonsohn’s two lines test to the median of each subject’s *K*-statistic and Lempel-Ziv complexity over all trials from their altered states (seizure, anesthesia, LSD), normalized to their own normal waking baseline (as shown in Fig. 3). The test failed to reject the null hypothesis of no inverse-U relationship between the normalized *K*-statistic and both univariate and concatenated Lempel-Ziv complexity, but not joint Lempel-Ziv complexity (Table 1). Moreover, as predicted, our within-subject analyses showed significant increases in chaoticity coinciding with significant drops in Lempel-Ziv complexity in the anesthesia state; small but significant reductions in chaoticity coinciding with significant increases in Lempel-Ziv complexity in the LSD state; and significant reductions in both chaoticity and Lempel-Ziv complexity during generalized seizures (SI Appendix, Fig. S4). Furthermore, we observed that the degree of reduction in chaoticity during the LSD state relative to placebo (assessed by normalizing each subject’s median *K*-statistic during their LSD state by their median during their normal waking state, as in Fig. 3) was significantly correlated with subjects’ behavioral ratings (see Materials and Methods) of the intensity of the LSD experience (partial correlation *ρ*=0.55,p=0.033, controlling for differences between placebo and LSD states in the median frequency at which signals were low-pass filtered). In order to assess whether median estimated chaoticity varied significantly across brain states, independently of the frequency at which signals were low-pass filtered, we performed a cross-subject non-parametric (permutation-based, 1000 permutations) analysis of covariance (ANCOVA), with median *K*-statistic as the response variable, brain state (i.e. normal waking, generalized seizure, anesthesia, or LSD) as the group label, and median frequency at which signals were low-pass filtered as the covariate. We observed significant variation in estimated chaoticity across states (F=61.765,p=0.001) with no effect of either median low-pass filter frequency (F=0.116,p=0.752) or interaction between median low-pass filter frequency and estimated chaoticity (F=0.214,p=0.959). The same result was obtained for chaoticity estimates normalized to each subject’s individual normal waking baseline (as reported in Fig. 3) (F=130.202,p=0.001), again with no effect of either median low-pass filter frequency (F=0.188,p=0.661) or interaction between median low-pass filter frequency and estimated chaoticity (F=0.414,p=0.922). Furthermore, our analyses of surrogate time-series not only suggest that low-frequency cortical electrodynamics are predominantly deterministic, but also show no difference in the level of stochasticity of cortical dynamics across brain states (SI Appendix, Tables S1-S2), which suggests that these between-condition differences were likely driven by changes in the relative stability of cortical dynamics across different brain states as predicted, rather than to changing levels of intrinsic noise in cortical networks. Finally, we compared the low-frequency power spectral densities of our real and simulated cortical electrodynamics, and observed spectral changes that were consistent across our real and simulated data (SI Appendix, Figs. S1-S3), which lends further support to the model’s prediction of increased or decreased chaoticity relative to the edge-of-chaos critical point in these different states.

**Fig. 3.**
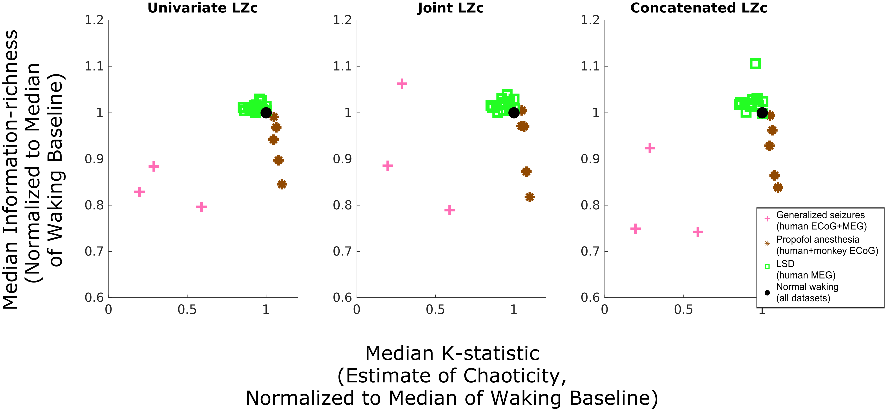
Transitions of low-frequency cortical electrodynamics away from the edge-of-chaos critical point induce a loss of information in cortical dynamics during unconscious states. We applied the modified 0-1 chaos test to ECoG and MEG recordings from humans and macaques across different brain states in order to empirically assess the predicted relationship between proximity to edge-of-chaos criticality, consciousness, and the information-richness of cortical dynamics. Here, each marker represents the median estimated chaoticity and information-richness of cortical dynamics across each individual subject’s trials, normalized to the median of their normal waking baseline. The observed inverse-U relationship between stability and information-richness, with cortical dynamics during conscious states at the top of this inverse-U, validates the prediction that cortical dynamics operate near the edge-of-chaos critical point during conscious states, transition deeper into the chaotic/unstable phase under GABAergic anesthesia, and transition into the periodic/stable phase during generalized seizures. These results support our hypothesis that these transitions away from edge-of-chaos criticality during unconscious states induce a loss of information in electrical cortical activity. Moreover, the counter-intuitive reduction of chaoticity coinciding with increased information-richness in the LSD state supports our prediction that waking cortical dynamics operate on the chaotic side of this critical point. See SI Appendix, Fig. S4 for statistical analysis of within-subject results.

### Edge-of-chaos criticality is a potential clinical biomarker of consciousness

The above findings support the hypothesis that the low-frequency electrodynamics of the cortex during conscious states are poised near the edge-of-chaos critical point, and specifically operate on the unstable side of this critical point. This implies that use of the modified 0-1 chaos test to assess the proximity of cortical electrodynamics to edge-of-chaos criticality, or to the unstable side of this phase transition, may be clinically useful as a novel tool for monitoring depth of anesthesia or diagnosing and monitoring emergence from disorders of consciousness - a group of conditions for which new biomarkers are sorely needed (45). Toward that end, we here introduce a novel time-series estimate *c* of proximity to edge-of-chaos criticality, based on a nonlinear transformation of the *K*-statistic (see Materials and Methods). Our measure *c* includes a parameter *α*, set between 0 and 1, such that *c* will approach 1 as the *K*-statistic approaches *α*, and will approach 0 as the *K*-statistic approaches either 0 (periodicity) or 1 (strong chaos). Note that *α* values nearer to 0 will bias our criticality measure to assign higher values to systems on the stable side of the edge-of-chaos critical point, while *α* values nearer to 1 will bias our measure to assign higher values to systems on the chaotic side of the critical point.

To test the diagnostic utility of this new criticality measure *c*, we applied our chaos analysis pipeline (i.e. low-pass filtering at a frequency determined by the FOOOF algorithm followed by application of the modified 0-1 chaos test) to clinical EEG data recorded from four traumatic brain injury patients as they recovered consciousness (see Materials and Methods). Degree of consciousness was assessed using the Glasgow Coma Scale (GCS) as part of conventional bedside neurobehavioral testing. Following prior work (46, 47), data were split into conscious and unconscious states based on the verbal and motor sub-scores of the GCS. Patients were considered conscious if either their GCS verbal sub-score was greater than or equal to four (meaning that they could answer questions) or if their motor sub-score was greater than or equal to five (meaning that they displayed clearly purposeful movement). We considered patients unconscious if their verbal sub-score was less than four and motor sub-score was less than five, though we note that this criterion cannot differentiate between unconsciousness and unresponsiveness/disconnectedness.

To test the utility of our criticality measure as a biomarker of consciousness, we converted the median *K*-statistics of these four patients in their unconscious and conscious states, along with the median *K*-statistics of our five anesthesia subjects and three generalized seizure subjects in their waking and unconscious states, to our new criticality estimate *c*, using 19 unique values of its parameter *α* ranging from 0.05 to 0.95 in steps of 0.05. For each value of *α*, we performed a cross-subject, right-tailed Wilcoxon rank-sum test to compare estimates of proximity to edge-of-chaos criticality in conscious versus unconscious states. Before correcting for multiple comparisons, estimates of criticality were significantly higher during conscious states for all *α* values between 0.65 and 0.85; after conservative Bonferroni-correction, *c* at *α*=0.85 remained significantly higher across subjects during conscious states than during unconscious states (p< 10^−4^ before Bonferroni correction, p=0.016 after Bonferroni correction) (SI Appendix, Fig. S5) (Fig. 4A). A cross-subject Wilcoxon rank-sum test revealed no significant difference in the median low-pass filter frequencies selected by the FOOOF algorithm in conscious vs unconscious states (p=0.795), while right-tailed Wilcoxon rank-sum tests showed that, across subjects, consciousness corresponded to significantly higher values of univariate Lempel-Ziv complexity (p=0.003) (Fig. 4B) and concatenated Lempel-Ziv complexity (p=0.0265) (SI Appendix, Fig. S6) but not joint Lempel-Ziv complexity (p=0.107) (SI Appendix, Fig. S6). Furthermore, after controlling for the median frequency at which signals were low-pass filtered across these twelve subjects (four DOC patients, five anesthesia subjects, and three generalized seizure subjects), our criticality measure *c* (at *α*=0.85) was significantly correlated with cross-trial median univariate Lempel-Ziv complexity (partial correlation *ρ*=0.66, p< 10^−4^) (Fig. 4C) and concatenated Lempel-Ziv complexity (*ρ*=0.66, p< 10^−4^) but not with joint Lempel-Ziv complexity (*ρ*=0.36, p=0.093) (SI Appendix, Fig. S6); these correlations support the hypothesis that proximity to the edge-of-chaos critical point mediates the information-richness of cortical electrodynamics as well as consciousness. Finally, we used a one-tailed block bootstrap test (block size = 30 seconds of data), which controls for the non-independence of successive time-points by preserving local time-series autocorrelations, to test for within-subject increases in *c* as patients recovered consciousness. We found significant increases in *c* for all four DOC patients (Fig. 4D), which supports the potential diagnostic utility of this new criticality measure. Significant within-subject increases in univariate Lempel-Ziv complexity were also observed within all four DOC patients as they regained consciousness, but not in joint or concatenated Lempel-Ziv complexity (SI Appendix, Fig. S7).

**Fig. 4.**
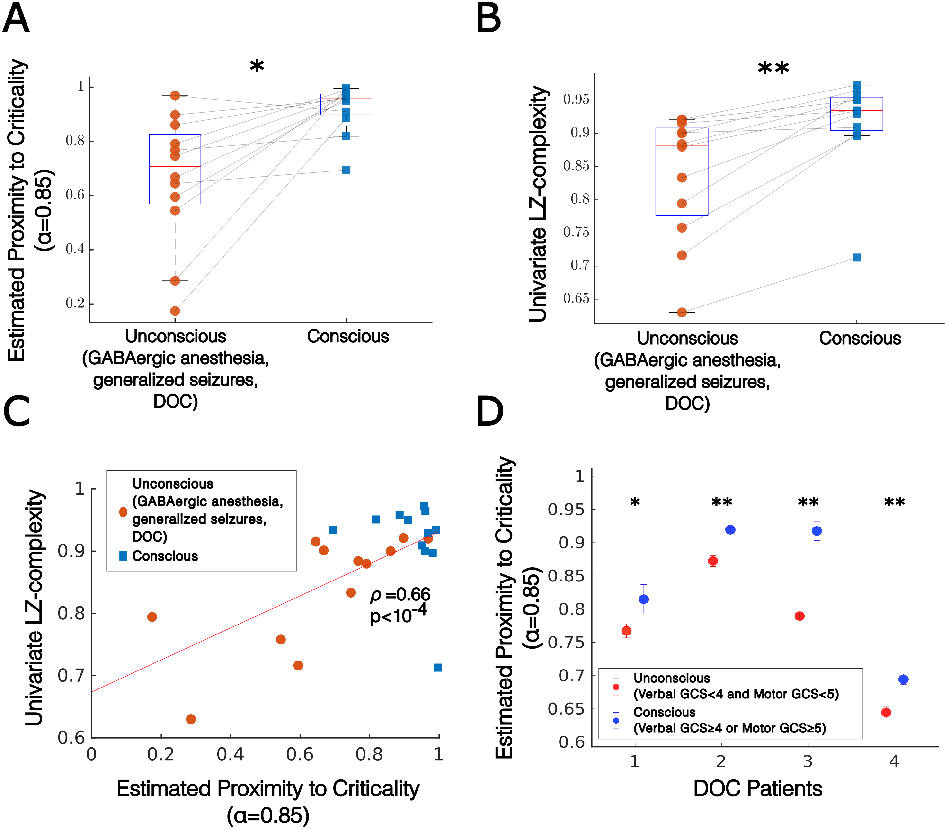
Criticality predicts consciousness. **A** Using our new time-series measure of criticality (derived from the 0-1 chaos test - see Materials and Methods), we estimated the proximity of low-frequency cortical dynamics to edge-of-chaos criticality in 12 subjects for whom data were available from both conscious and unconscious states (namely, five GABAergic anesthesia subjects, three generalized seizure subjects, and four DOC patients). Our criticality measure includes a parameter *α*, which we here set to 0.85, based on our parameter analysis (see SI Appendix, Fig. S5). Estimates of proximity to edge-of-chaos criticality were significantly higher (p< 10^*−*4^ before Bonferroni correction for comparisons at multiple values of *α*, and p=0.0157 after Bonferroni correction) in conscious states than in unconscious states (significance was tested using a right-tailed Wilcoxon rank-sum test). **B** Cross-trial, within-subject medians of univariate Lempel-Ziv complexity were significantly higher (p=0.003) during conscious states than during unconscious states. See SI Appendix, Fig. S6 for comparisons using joint and concatenated Lempel-Ziv complexity. **C** Across the waking (blue square) and non-waking (red circle) states of all 12 subjects exhibiting transitions between consciousness and unconsciousness, cross-trial medians of estimated proximity to edge-of-chaos criticality (with *α*=0.85) were significantly correlated with cross-trial medians of univariate Lempel-Ziv complexity (partial correlation *ρ*=0.66, p< 10^*−*4^, controlling for median frequency at which signals were low-pass filtered). See SI Appendix, Fig. S6 for comparisons using joint and concatenated Lempel-Ziv complexity. **D** As was the case for our cross-subject analysis (**A**), our within-subject, cross-trial analysis revealed significant increases in our criticality measure (with *α*=0.85) in four DOC patients as they recovered consciousness. Significance was assessed using a left-tailed overlapping block bootstrap test (which controls for dependencies across data points by preserving local time-series autocorrelations) with a block size of three trials (30 seconds of recording), to test against the null hypothesis that median estimated proximity to criticality during conscious states is not greater than median estimated proximity to criticality during unconscious states. Circles correspond to cross-trial medians, and errorbars indicate standard error of the median (estimated by taking the standard deviation of a bootstrap distribution of sample medians) * p<0.05, ** p<0.01.

## Discussion

In this paper, we present the first empirical evidence that cortical electrodynamics exhibit a high information-carrying capacity during conscious states by operating near the mathematically specific critical point separating periodicity and chaos. Our evidence was based on the first application (to our knowledge) of the recently developed modified 0-1 chaos test to neural electrophysiology data. Many systems, including deep neural networks (24), have been shown to exhibit their highest information-processing capacity when poised near this transition from periodicity/stability to chaos/instability (6–9, 34), likely because dynamics near this critical point optimally balance stability with flexibility and responsiveness to inputs (48). Both our simulation and empirical results suggest that waking cortical dynamics specifically operate on the chaotic/unstable side of this phase transition, which supports the decades-old conjecture that the waking brain might utilize weak dynamical chaos in the service of efficient information processing (49), particularly during conscious states (21). From a computational perspective, it is reasonable that evolution would have tuned waking, conscious cortical dynamics to the chaotic side of this critical point, because traversing this critical point into the chaotic phase coincides with a transition from narrow-band to broadband, multi-frequency oscillations (50), a phenomenon which has been exploited in the engineering context to enable frequency multiplexing (i.e., carrying of information at multiple frequencies) (51); tellingly, such frequency multiplexing is thought to be ubiquitous in mammalian neurodynamics during normal waking states (52). Note that this finding that cortical electrodynamics operate on the chaotic side of criticality during normal waking states is fully consistent with the hypothesis that conscious cortical electrodynamics operate on the “ordered” side of criticality, because, as mentioned in the Introduction, chaos is in fact the “ordered” phase of a dynamical system with respect to this critical point (21) (see SI Appendix, Supplementary Note 1). This result is also consistent with findings that cortical dynamics operate near the critical onset of neuronal avalanches; this is because the neuronal avalanche critical point is distinct from the edge-of-chaos critical point and likely occurs *within* the weakly chaotic regime of neural networks (17), precisely where our results suggest cortical dynamics lie during normal conscious states.

We further present evidence that transitions of cortical electrodynamics away from the edge-of-chaos critical point - either deeper into the chaotic/unstable phase, as our evidence suggests is the case for GABAergic anesthesia, or into the periodic/stable phase, as our evidence suggests is the case for generalized seizures - precipitate a loss of information-richness in cortical dynamics and unconsciousness. These results are consistent with previous findings of a loss of empirical signatures of criticality during these states of unconsciousness (4, 13, 14), but go beyond prior analyses in specifying whether dynamics in these states are sub-critical or super-critical with respect to a specific, mathematically well-defined critical point (in this case, the edge-of-chaos critical point). Finally, we present evidence that psychedelics may increase the information-richness of cortical electrodynamics by moderately stabilizing cortical activity, i.e., by approaching the edge-of-chaos critical point from the chaotic/unstable side of the edge. This result not only supports prior findings suggesting a transition c loser to criticality in the LSD state (44), but also confirms the model-based prediction that normal waking cortical dynamics specifically operate on the unstable side of the edge-of-chaos critical point.

We note that our finding o f i ncreased i nstability during GABAergic anesthesia may appear to conflict w ith a prior report by Solovey and colleagues of increased stability in the cortical dynamics of macaques during propofol anesthesia (53). This seeming discrepancy rests on differing notions of stability, as well as different assumptions about data: Solovey and colleagues defined stability in terms of the eigenvalues of regression matrices estimated from ECoG recordings, a notion of stability which only indicates that a process will not diverge to infinity, and which further assumes that data are both linear and stochastic (an assumption not supported by our analyses - see SI Appendix, Tables S1-S2). In contrast, we assessed stability in terms of sensitivity to perturbations/inputs, and also used time-series analysis tools which do not assume linearity, and which therefore capture features of data that cannot by definition be captured by linear analysis tools such as autoregressive models. It is also worth noting that two out of the four ECoG data sets used in the report by Solovey and colleagues were the same as the macaque anesthesia data used here (data were downloaded from the same repository - see Materials and Methods), and yet we found robust increases in instability in the anesthetized state for these two macaques, as we did in our three human anesthesia subjects (Fig. 2, SI Appendix, Fig. S4). While the finding that GABAergic anesthetics destabilize cortical electrodynamics may be counter-intuitive, this possibility is further suggested by prior observations of disrupted long-range cortical phase coherence during propofol anesthesia, which is a key prediction of this anesthesia-as-chaos mean-field model (54).

We note that although our criticality measure *c* increased in all four DOC patients as they regained consciousness, estimates of chaoticity were significantly higher (within-subject) during unconsciousness in only three out of four of the patients (similar to the GABAergic anesthesia state) and were significantly lower during unconsciousness in the fourth patient (similar to generalized seizures) (SI Appendix, Fig. S7). This may imply that disorders of consciousness constitute a heterogeneous set of conditions with respect to the stability of cortical electrodynamics, a possibility we hope to explore more fully in future work. We further note one important limitation in our analysis of DOC patients, which is the potential confounding effect of drugs administered to the patients: patients were occasionally administered several painkillers and anesthetics on the same day as GCS assessments and EEG data collection (SI Appendix, Table S5) (Materials and Methods). We were unable to ascertain the precise timing of drug administration relative to behavioral assessments and, as such, we cannot rule out the possibility that observed differences in cortical stability/criticality in unconscious states versus conscious states in these DOC patients were possibly driven by the effects of these drugs on their cortical electrodynamics. Moreover, our sample size of DOC patients who regained consciousness was small (n=4), and so the utility of our criticality measure *c* as a biomarker of consciousness in patients with disorders of consciousness warrants validation in a larger dataset. Along the same lines, if this framework is to be used in the aid of diagnosis, then it will be imperative to develop additional methods for estimating changing levels of chaoticity in cortical electrodynamics. This might be achieved, for example, by observing the consistency of cortical responses to external stimuli (e.g. in response to transcranial magnetic stimulation) - a possibility we plan to explore in future work.

Finally, we note that it would be fruitful to further study neural computation near the edge-of-chaos critical point on a more theoretical level. While important advances have been made along these lines, for example in establishing relationships between this critical point and the trainability of deep neural networks (24), information complexity (6–9), Bayes-optimal perceptual categorization (55), and combinatorial optimization (56), much theoretical work remains to be done to understand the implications of these findings for neural computation. If the electrodynamics of the cortex during conscious states operate near this critical point, as our work suggests, then improving our theoretical understanding of computation at the onset of chaos will also improve our understanding of how, precisely, neural computation is disrupted in unconsciousness.

## Materials and Methods

### Mean-Field Model Equations of Cortical Electrodynamics

We here study the mean-field model of Steyn-Ross, Steyn-Ross, and Sleigh (26). The model allows for straightforward manipulation of both the strength and balance of postsynaptic inhibition and excitation, which have long been thought to be key in tuning neural dynamics to chaotic (57), critical (58), and information-rich (58) states. Furthermore, the model is unique in its inclusion of junction coupling between inhibitory interneurons, which recent empirical work in zebrafish has shown are also likely important for tuning neural dynamics toward and away from criticality (30).

The model simulates GABAergic anesthesia (e.g. propofol or sevoflurane) as an increase in cortical inhibition coupled with a mild decrease in gap junction coupling between inhibitory interneurons, based on findings that GABAa agonists (59), and GABAergic anesthetics more specifically (60), inhibit gap junction communication (59, 60), and that these compounds also increase postsynaptic inhibition by prolonging inhibitory postsynaptic potentials (61). The model treats waking conscious states as a balance between excitation and inhibition, with strong gap junction coupling between inhibitory interneurons, which yields weak chaos (near edge-of-chaos criticality) in the model’s deterministic component (Fig. 2A), arising from interacting Turing (spatial) and Hopf (temporal) instabilities. Finally, a strong reduction of inhibitory gap junction coupling results in a Hopf bifurcation that produces periodic dynamics reminiscent of whole-of-cortex, generalized seizures (26). This is consistent with observations of increased seizure frequency following either genetic ablation (62) or drug-induced reduction (63) of gap junction coupling between inhibitory interneurons. See Steyn-Ross, Steyn-Ross, and Sleigh (26) for full details on model parameters.

The mean excitatory and inhibitory potentials *V*_*e*_ and *V*_*i*_ of each simulated neural population in the mean-field model, positioned at a location 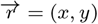, are described by:

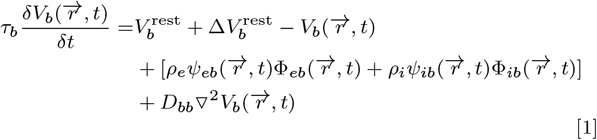

where presynaptic to postsynaptic directionality is indicated by the right arrow, the subscript *e* indicates a presynaptic excitatory neural population, the subscript *i* indicates a presynaptic inhibitory neural population, and the subscript *b* indicates either a postsynaptic excitatory or postsynaptic inhibitory neural population. The bracketed term in Eq. 1 represents voltage inputs via chemical synapses, and the final term in Eq. 1 represents voltage inputs from diffusive gap junction coupling. ▽^2^ is the 2D Laplacian operator. *D*_*bb*_ represents the strength of diffusive gap junction coupling between adjacent neurons, such that *D*_*ee*_ is gap junction coupling between excitatory populations and *D*_*ii*_ is gap junction coupling between inhibitory populations. Because there is far more abundant gap-junction coupling between inhibitory interneurons than excitatory neurons (64), *D*_*ee*_ is set to *D*_*ii*_/100. *D*_*ii*_ is one of the key biological parameters we vary. For a given excitatory or inhibitory neural population, 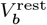 is the mean resting potential, *τ*_*b*_ is the soma time constant, and *ρ*_*b*_ is the strength of chemical synapse coupling, which is scaled by the following reversal-potential function *ψ*_*ab*_:

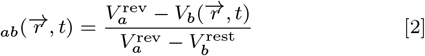

which equals one when a neuron is at its resting potential and equals 0 when the membrane potential equals the reversal potential. For excitatory AMPA receptors, 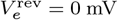, and for inhibitory GABA receptors, 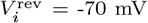. The Φ_*ab*_ functions in Eq. 1 describe postsynaptic*i* spike-rate fluxes:

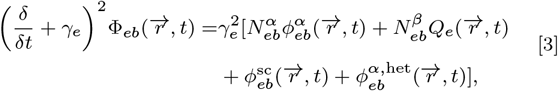

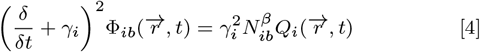

where the *α* superscript corresponds to inputs from long-range myelinated axons: 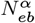 is the number of axonal inputs to a population and 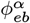 is long-range spike-rate flux. The *β* superscript corresponds to inputs from short-range chemical synapses, such that 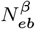 is the number of local chemical synapses in a neural population. *Q*_*e,i*_ is the local spike-rate flux, and 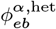 is a heterogeneous flux input. 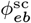 is white noise, taken to represent random inputs to the cortex from subcortical sources (e.g. sensory inputs); note that the inclusion of a noise term means that the above equations are stochastic differential equations, and that analyses of the ground-truth chaoticity of the model (i.e. its largest Lyapunov exponent) are performed exclusively using the non-stochastic components of the model equations; estimates of chaoticity using the 0-1 test (see below) are performed with the model’s noise input turned on, so as to better assess the viability of detecting changing levels of chaoticity in real cortical recordings. *γ*_*i*_ is the inhibitory rate constant and *γ*_*e*_ is the excitatory rate constant, which we vary so as to the study the effect of excitation and inhibition on chaos in the model. See Steyn-Ross, Steyn-Ross, and Sleigh (26) for more details on the model equations. Other than the inhibitory gap-junction coupling strength *D*_*ii*_, the excitatory rate constant *γ*_*e*_, and the inhibitory rate constant *γ*_*i*_ (all of which we vary in our parameter sweep), all parameters in our simulations are unchanged from the original model, and are taken from the empirical literature (26). *D*_*ii*_ was varied from 0.1 to 0.7 in steps of 0.2, and both *γ*_*e*_ and *γ*_*i*_ were varied from 0.945 to 1.05 in steps of 0.005. Of the 1,936 resulting simulations, 1,160 yielded flat, non-oscillatory activity, likely reflecting stable fixed points of the model; these fixed point solutions were excluded from all analyses, because these non-oscillatory solutions would likely yield high estimates of Lempel-Ziv complexity simply due to the information-richness of the noise perturbations rather than of the underlying system dynamics. This left 776 unique model simulations of oscillatory behavior. Based on prior work (26), the waking conscious simulation corresponded to *γ*_*e*_ = 1, *γ*_*i*_ = 1, and *D*_*ii*_ = 0.7. The anesthesia simulation corresponded to *γ*_*e*_ = 1, *γ*_*i*_ = 1.015, and *D*_*ii*_ = 0.5, and the seizure simulation corresponded to *γ*_*e*_ = 1, *γ*_*i*_ = 1, and *D*_*ii*_ = 0.1. The nearest-to-criticality and maximally information-rich state of the model (see SI Appendix, Figs. S1, S3) corresponded to *γ*_*e*_ = 1.04, *γ*_*i*_ = 1, and *D*_*ii*_ = 0.5. The model equations were integrated using a forward time center spaced first-order Euler method, with an integration step of 0.2 ms. Simulated electrodynamics were then downsampled to a sampling frequency of 500 Hz, and the final 10 seconds (i.e. 5,000 time-points) were extracted from the downsampled data, so as to perfectly match the length and sampling frequency of the ECoG and MEG datasets analyzed in this paper.

### Lempel-Ziv Complexity

Lempel-Ziv complexity is a measure of the size of a signal following Lempel-Ziv compression, and thus tracks the amount of non-redundant information in a signal (31). To compute Lempel-Ziv complexity, a continuous recording must first be discretized. Following prior work (42, 65), we binarized both our simulated and recorded data by thresholding at the mean of the signal’s instantaneous amplitude, which is the absolute value of the analytic signal; the analytic signal is 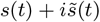, where *s*(*t*) is the original time-series signal, *i* is the imaginary unit, and 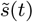 is the Hilbert transform of *s*(*t*). We then computed three measures of Lempel-Ziv complexity: 1) the median univariate Lempel-Ziv complexity across all recorded channels (“Univariate LZc”), 2) the joint Lempel-Ziv complexity between all channels, using the method described by Zozor and colleagues (66), and 3) the Lempel-Ziv complexity of all channels concatenated, time-point by time-point, into a single string, following the method described by Schartner and colleagues (42, 65). Typically, Lempel-Ziv complexity is then normalized to provide a single value between 0 and 1. We compared several different normalization approaches, and found that the approach most robust against changes to a signal’s spectral profile was to divide the Lempel-Ziv complexity of a signal by the Lempel-Ziv complexity of a phase-randomized surrogate of that signal (SI Appendix, Fig. S8), following Brito and colleagues (67); note that phase-randomized surrogates were generated independently for each channel-x-trial in all recordings for the calculation of the Lempel-Ziv complexity measures. All measures of Lempel-Ziv complexity reported in this paper were normalized in this fashion, and were calculated for data low-pass filtered at 45 Hz. Data were low-pass filtered at 45 Hz to avoid potential confounds introduced by muscle activity at higher frequencies.

### Calculating Largest Lyapunov Exponents in the Mean-Field Model

The ground-truth chaoticity of a system is determined by its largest Lyapunov exponent, which is the rate of divergence between initially similar trajectories in a system’s phase space. A positive largest Lyapunov exponent means that a system is chaotic, because it indicates exponential divergence of initially similar system states. A negative largest Lyapunov exponent indicates periodicity, because it indicates exponentially fast convergence of initially similar states. A largest Lyapunov exponent near 0 corresponds to edge-of-chaos criticality, and near-0 exponents indicate that a system is near the edge-of-chaos critical point. The larger the largest Lyapunov exponent, the more strongly chaotic the system is. Following Steyn-Ross, Steyn-Ross, and Sleigh (26), we estimate the largest Lyapunov exponent of the mean-field model by simulating two runs of its deterministic component (i.e., with its noise inputs turned off), with slightly different initial conditions. The divergence between the excitatory firing rate of run 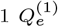 and run 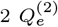 is estimated as their summed squared-difference *ϵ(t)* down the midline of the simulated cortical grid:

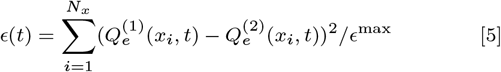

where *ϵ*^max^ is a normalization parameter, which equals the maximum possible difference between the two runs:

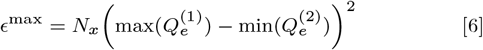

where *N*_*x*_=120, i.e. the number of simulated neural populations in the cortical sheet. The rate of divergence between the two runs *ϵ*(*t*) is directly related to the largest Lyapunov exponent Λ of the system:

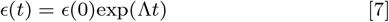

where *ϵ*(0) is the distance between the two runs at *t* = 0. The largest Lyapunov exponent can therefore be estimated by measuring the slope of ln*E*(*t*)-versus-*t*. A positive slope indicates a positive largest Lyapunov exponent (and therefore chaotic dynamics), a negative slope indicates periodicity, and a flat slope indicates edge-of-chaos criticality.

### Extracting Low-Frequency Cortical Activity

The mean-field model described above specifically simulates the low-frequency (<4 Hz) component of macro-scale electric cortical oscillations. To compare the model results against real data, we therefore extracted the low-frequency component of both our simulated and real cortical signals. Although different frequencies of cortical electrodynamics have historically been studied at fixed, canonical frequency bands, with choices of oscillation center frequencies and bandwidths varying across studies, there is growing evidence that these center frequencies and bandwidths vary considerably as a function of age, brain state, subject, and species, and that low-pass filtering at fixed canonical frequencies can therefore produce spurious oscillations where no oscillations exist (41). Given that our analyses span diverse brain states, species, and imaging modalities, it was important to identify subject-, trial-, and channel-specific neural oscillation frequencies. We therefore identified low-frequency neural activity for each channel, for each trial, using the recently developed “Fitting Oscillations and One Over F” or “FOOOF” algorithm, which automatically parameterizes neural signals’ power spectra (41). The algorithm fits a neural power spectrum as a linear combination of the background 1/f component with oscillations at specific frequencies that rise above this background 1/f component as peaks in the power spectrum. The algorithm fits the spectral power *P* as:

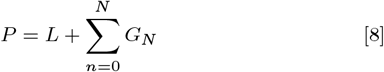

where *L* is the background 1/f power spectrum, and each *G*_*n*_ is a Gaussian fit to a peak rising above the 1/f background:

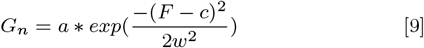

where *a* is a given oscillation’s amplitude, *c* is its center frequency, *w* is its bandwidth, and *F* is a vector of input frequencies. The 1/f background component *L* is modeled as an exponential in semilog-power space (i.e. with log power values as a function of linear frequencies):

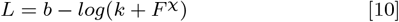

where *b* is a broadband power offset, *χ* is the spectral slope, *k* controls the “knee” at which the 1/f power spectrum bends, and *F* is a vector of input frequencies.

To specifically extract the low-frequency component of neural oscillations, we set the input frequency range *F* to 1-6 Hz. The FOOOF algorithm then identifies the center frequencies and bandwidths of putative oscillations that rise above the 1/f background within this frequency range. For all channels-x-trials in our data, we extracted the lowest frequency oscillation identified by the algorithm, by low-pass filtering at the high-frequency end of the bandwidth of the slowest identified oscillation. If the FOOOF algorithm failed to identify an oscillation in the 1-6 Hz range for a particular channel in a particular trial, then data for that channel in that trial were excluded from further analysis. Across all datasets, the mean frequency selected using this approach was 3.27 Hz, with a standard deviation of 0.48 Hz. We then low-pass filtered all signals using EEGLAB’s two-way least-squares FIR low-pass filtering, where the filter order was set to 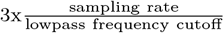 (the default of EEGLAB). Note that using the FOOOF algorithm improved our ability to track chaoticity in the mean-field model of cortical electro-dynamics, where the ground truth chaoticity is known (SI Appendix, Tables S3-S4), and that estimates of the chaoticity of data low-pass filtered using the FOOOF algorithm were stable across different simulations (SI Appendix, Fig. S9), which validates its utility in tracking chaoticity in real low-frequency cortical electrodynamics.

### The Modified 0-1 Test for Chaos

The 0-1 chaos test was developed by Gottwald and Melbourne (36) as a simple tool for testing whether a discrete-time system is chaotic, using only a single time-series recorded from that system. Gottwald and Melbourne provided an early modification to the test, which made it more robust against measurement noise (37). Dawes and Freeland added additional modifications to the test, improving its ability to distinguish between chaotic dynamics and strange non-chaotic or quasiperiodic dynamics (39). The modified 0-1 test takes a univariate time-series **ϕ**, and uses that time-series to drive the following two-dimensional system:

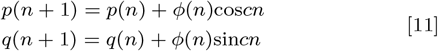

where *c* is a uniformly distributed random variable bounded between 0 and 2*π*. For a given *c*, the solution to Eq. 1 yields:

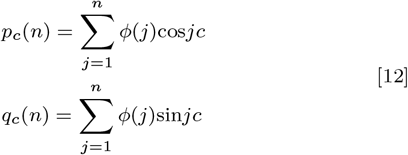

If the time-series **ϕ**is periodic, the motion of **p** and **q** is bounded, while if the time-series **ϕ** is chaotic, **p** and **q** display asymptotic Brownian motion. The time-averaged mean square displacement of **p** and **q** is

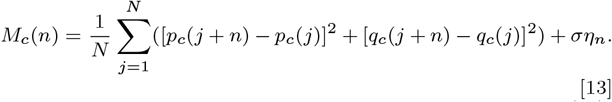

where *η*_*n*_ is a uniformly distributed random variable between 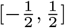 and *σ* controls the amplitude of the added random variable *η*_*n*_. We set *σ* to 0.5 and normalized the standard deviation of all signals to 0.5, based on our previously published analyses (25) of the effect of different parameter values for 0-1 test performance across diverse datasets. To compute the degree of chaos using a single statistic *K*, the 0-1 test calculates the growth rate of the mean squared displacement of the two-dimensional system in Eq. 5 using a correlation coefficient:

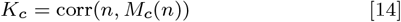

*K* is computed for 100 different values of *c*, uniformly randomly sampled between 0 and 2*π*, and the output of the test is the median *K* across different values of *c*. As the length of a time-series is increased, this median *K* value will approach 1 for chaotic systems, and will approach 0 for periodic systems, and will track degree of chaos for finite-length time-series (36–39). For both our real and simulated cortical activity, we calculated *K* for every channel in a trial, and estimated that trial’s level of chaoticity as the median *K*-statistic across all channels in that trial.

The 0-1 test is designed to detect and track chaos in discrete-time systems, and thus signals recorded from non-time-discrete processes (like neural electrodynamics) must first be transformed into discrete-time signals before application of the test (38). Two approaches have been proposed for signal time-discretization prior to application of the test: downsampling (38), or taking all local minima and maxima of a continuous signal (40). We here used the latter approach for all datasets (real and simulated), as it yielded best correspondence to the ground-truth in our mean-field simulations (SI Appendix, Tables S3-S4).

### A New Time-Series Estimate of Proximity to Edge-of-Chaos Criticality

With an eye toward clinical applications of this edge-of-chaos criticality framework in the study of unconsciousness, we here introduce a new time-series estimate of proximity to the edge-of-chaos critical point, based on the *K*-statistic outputted by the modified 0-1 chaos test (see above). This new measure *c* is defined as follows:

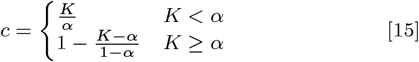

where *K* is the output of the 0-1 chaos test and *α* is a parameter that takes on a value between 0 and 1. This criticality measure *c* will approach 1 as *K* approaches *α*, and will approach 0 as *K* approaches either 0 (periodicity) or 1 (strong chaos). As noted in the Results, precise choice of *α* may bias *c* toward either periodic near-critical or chaotic near-critical dynamics (i.e., to dynamics on either the stable or unstable side of the edge-of-chaos critical point), and thus the optimal value of *α* for potential clinical assessments of consciousness using *c* will need to be determined by further empirical work.

### Epilepsy Data

Surface ECoG data from nine epilepsy patients were downloaded from the European Epilepsy Database (68). Of these, only two subjects experienced fully generalized seizures (in both cases, seizures were focal with secondary generalizations). Subject 1 was a 42 year old male with epilepsy caused by right cortical dysplasia, and who was receiving the anticonvulsant medication lamotrigine. The subject had six intracranial electrode strips (26 electrodes in total) placed over right lateral temporal cortex to monitor seizure focus. Subject 2 was a 14 year old female with cryptogenic epilepsy (i.e. unknown cause) who was receiving the anticonvulsant medications valproate and topiramate; the subject had one grid and six electrode strips (96 electrodes total) placed over left temporal and lateral left temporal cortex to monitor seizure focus. Signals from both subjects were recorded at a sampling rate of 1024 Hz. Data were demeaned, detrended, and bandstop filtered at 50 Hz and harmonics (the line noise frequency in Europe). Data were resampled to 500 Hz, divided into 10-second trials, and re-referenced to the common average. For the seizure state, we only included trials for which seizures were fully generalized across all channels for the entire trial duration. The data were then visually inspected for artifacts. Data from electrodes with consistent motion or drift artifacts were removed, and 10-second trials with large motion artifacts spanning multiple electrodes were removed.

Additionally, a magnetoencephalograpy (MEG) recording of one patient’s generalized absence seizure, previously published by Dominguez and colleagues (69), was re-analyzed. Data were provided by D.M.M. Note that MEG datasets were recorded for two other epilepsy patients by Dominguez and colleagues, but that these were for tonic seizures; the muscle convulsions during these tonic episodes produced large motion artifacts in the MEG data, which rendered analysis of low-frequency periodicity impossible. These datasets were therefore not analyzed. The patient whose data were re-analyzed in the present paper (Seizure Subject 3 in SI Appendix, Figs. S1-2) was an 18 year-old female who was receiving a low dose of valproate, and with no reported structural abnormalities or prior brain surgery. Data from this patient were recorded at 625 Hz using a CTF Omega 151 channel whole-head system (CTF Systems, Port Coquitlam, British Columbia, Canada). Data were split into 10-second trials, demeaned, detrended, and bandstop filtered at 60 Hz and harmonics (the line noise frequency in Canada, where data were collected). Data were then visually inspected. Consistently motion or drift artifact-affected channels were removed, and trials with large motion artifacts across channels were removed. Data were then downsampled to 500 Hz. We then ran an independent components analysis on the data, and removed components that corresponded to ocular or cardiac artifacts.

See SI Appendix, Fig. S10 for 10-second time-traces of these subjects’ cortical electrodynamics during generalized seizures.

### Human Anesthesia Data

Surface ECoG recordings from three human epilepsy patients given propofol anesthesia prior to surgical resection of their epileptic focus were analyzed. Data were collected at the University of California at Irvine, Medical Center. All patients provided informed consent in accordance with the local ethics committees of the University of California, Irvine (University of California at Irvine Institutional Review Board Protocol Number 2014-1522) and University of California, Berkeley (University of California at Berkeley Committee for the Protection of Human Subjects Protocol Number 2010-01-520), and provided written consent before data recording. Electrode placement was determined only by clinical criteria (Ad-Tech, SEEG: 5 mm inter-electrode spacing; Integra, Grids: 1 cm, 5 or 4 mm spacing). ECoG data were recorded using a Nihon Kohden recording system (256 channel amplifier, model JE120A), analogue-filtered above 0.01 Hz and digitally sampled at 5 kHz.

Patient 1 (Human Anesthesia Subject 1 in SI Appendix, Figs. S1-2) was a right-handed 25 year-old female with a diffuse lesion in the right supplementary motor area. The patient had one 8×8 grid placed over the right frontal lobe, covering superior temporal gyrus, postcentral gyrus, inferior parietal lobule, superior temporal gyrus, precentral gyrus, middle frontal gyrus, inferior frontal gyrus, and middle temporal gyrus; and two 2×5 anterior interhemisphere bilateral grids and two 2×8 posterior interhemisphere bilateral grids covering superior frontal gyrus and medial frontal gyrus, for a total of 116 cortical contacts. See SI Appendix, Fig. S11 for MRI scans with Patient 1’s cortical grids. The patient received 100 mg of propofol and 100 mcg of fentanyl prior to surgical resection of their epileptic focus. Their “waking conscious” data consisted of the twenty minutes prior to anesthetic induction, and their “anesthesia” data consisted of the twenty minutes following the loss of responsiveness to verbal commands.

Patient 2 (Human Anesthesia Subject 2 in SI Appendix, Figs. S1-2) was a 46 year-old, right-handed female with a lesion in the left supplemantary motor area. The patient had one 8×8 grid placed over the left frontal lobe, covering middle temporal gyrus, superior temporal gyrus, inferior frontal gyrus, middle frontal gyrus, superior frontal gyrus, precentral gyrus, superior temporal gyrus, postcentral gyrus, and inferior parietal lobule; one 2×8 strip placed over left medial cortex, covering left medial frontal gyrus, left cingulate gyrus, and left superior frontal gyrus; and one 2×8 strip placed over right medial cortex, covering right medial frontal gyrus, right cingulate gyrus, and right superior frontal gyrus, for a total of 96 contacts. See SI Appendix, Fig. S11 for MRI scans with Patient 2’s cortical grids. The patient received 140 mg of propofol and 50 mcg of fentanyl prior to surgery; at loss of consciousness, the patient received 50 mg of the muscle relaxant rocuronium; four minutes after loss of consciousness, the patient began receiving sevoflurane (another GABAergic anesthetic) for maintenance of anesthesia; note that the predictions of the mean-field model regarding the anesthesia state still hold for a combination of propofol and sevoflurane, as the model predictions should pertain to any GABAergic anesthetic. The patient’s “waking conscious” data consisted of the 19 minutes prior to anesthetic induction, and their “anesthesia” data consisted of the 16.8 minutes following the loss of responsiveness to verbal commands.

Patient 3 (Human Anesthesia Subject 3 in SI Appendix, Figs. S1-2) was a 20 year-old right-handed female who had previously received a left temporal lobectomy. The patient had one 4×8 grid placed over left frontal cortex, with contacts over middle frontal gyrus, Brodmann area 9, inferior frontal gyrus, superior temporal gyrus, middle temporal gyrus, precentral gyrus, and superior frontal gyrus; another 4×4 grid over left frontal cortex, with contacts over the orbital gyrus, inferior frontal gyrus, middle frontal gyrus, superior frontal gyrus, rectal gyrus, and superior temporal gyrus; a 2×6 grid over the temporal lobe, with contacts over the fusiform gyrus, inferior temporal gyrus, and middle temporal gyrus, as well as four contacts over the declive and one over the tuber of the cerebellum; and one 8×8 grid with contacts over parts of parietal, temporal, and occipital cortices, including postcentral gyrus, precentral gyrus, inferior parietal lobule, superior parietal lobule, Brodmann area 40, Brodmann area 7, supramarginal gyrus, and superior temporal gyrus, for a total of 124 surface electrodes. See SI Appendix, Fig. S11 for MRI scans with Patient 3’s ECoG grids. The patient received 150 mg of propofol and 100 mcg of fentanyl prior to surgical resection of their epileptic focus. Their “waking conscious” data consisted of the eight minutes prior to anesthetic induction, and their “anesthesia” data consisted of the 10.17 minutes following the loss of responsiveness to verbal commands.

Signals for all three patients were recorded at a sampling rate of 1,000 Hz. Epileptic activity was assessed by an experienced neurologist (R.T.K) and removed. Data were split into 10-second trials, demeaned, band-stop filtered at 60 Hz and harmonics (the line noise frequency in the United States, where data were collected), de-trended, downsampled to 500 Hz, and re-referenced to the common average. Data were then visually inspected for artifacts. Data from electrodes with consistent drift or motion artifacts were removed, and 10-second trials with large motion artifacts spanning multiple electrodes were removed.

### Macaque Anesthesia Data

Open-source ECoG recordings spanning the left cortices (including occipital, parietal, temporal, and frontal lobes) of two male macaques were downloaded from Neurotycho.org (70). See SI Appendix, Fig. S11 for an MRI scan showing the electrode placement of Macaques 1 and 2. Data were collected during awake/resting and propofol anesthesia states. The macaques were seated with head and arm movement restricted. Macaque 1 (Macaque Anesthesia Subject 1 in SI Appendix, Figs. S1-2) was intravenously administered 5.2 mg/kg of propofol, and Macaque 2 (Macaque Anesthesia Subject 2 in SI Appendix, Figs. S1-2) was intravenously administered 5 mg/kg of propofol. Loss of consciousness was determined by the emergence of low-frequency oscillations and the cessation of responses to physical stimuli. All data for the propofol condition are from the macaques’ unconscious state, and all data from the awake condition are from the macaques’ eyes-open state (i.e., data for which the eyes were covered were excluded). Signals were recorded at a sampling rate of 1,000 Hz. Data were split into 10-second trials, demeaned, band-stop filtered at 50 Hz and harmonics (the line noise frequency in Japan, where data were collected), detrended, downsampled to 500 Hz, and re-referenced to the common average. Data were then visually inspected for motion artifacts. Data from electrodes with consistent artifacts were removed, and 10-second trials with artifacts spanning multiple electrodes were removed.

### Human Lysergic Acid Diethylamide Data

Previously published (71) MEG recordings of nineteen humans following intravenous administration of either 75 *μ*g of lysergic acid diethylamide (LSD) or a saline placebo were re-analyzed. These data were provided by S.M. and R.C. Data from three subjects were excluded because of persistent motion or drift artifacts in their MEG signal across most trials. Of the sixteen remaining subjects, three were females, and the average age was 32.06 (with a standard deviation of 7.71 years). Due to the slow pharmacodynamics of LSD, MEG data were recorded four hours after drug administration. Subjects lay in a supine position during data acquisition. MEG signals were recorded using a CTF 275-channel radial gradiometer system with a sampling frequency of 1200 Hz. After the MEG recordings were collected, visual analogue scale ratings of the intensity of the LSD experience (on a scale from 0 to 20 in increments of 1) were presented to subjects on a projection screen visible from inside the scanner, which the subjects completed via button press (see Carhart-Harrris et al (71) for more details). MEG data were split into 10-second trials, demeaned, detrended, and bandstop filtered at 50 Hz and harmonics (the line noise frequency in the United Kingdom, where data were collected). Data were then visually inspected. Consistently motion or drift artifact-affected channels were removed, and trials with large motion artifacts across channels were removed. Data were then downsampled to 500 Hz. We then ran an independent components analysis on the data, and removed components that corresponded to ocular or cardiac artifacts.

### Clinical DOC data

Data were collected from four traumatic brain injury (TBI) patients admitted at the UCLA Ronald Reagan University Medical Center intensive care unit (ICU). Several criteria were applied for participation in the study in order to limit the investigation to those patients recovering from unconsciousness. Inclusion criteria: Glasgow Coma Scale (GCS) score ≤ 8 or an admission GCS score of 9-14 with computed tomography (CT) evidence of intracranial bleeding. Exclusion criteria: GCS > 14 with non-significant head CT, history of neurological disease or TBI, and brain death. The UCLA institutional review board approved the study. Informed consent was obtained according to local regulations. To manage symptoms and/or reduce cerebral metabolism, medications were administered to patients as needed, noted on a daily basis and sorted into appropriate categories: propofol, barbiturates, benzodiazepines, opioids, and dissociative anesthetics. Behavioral assessments were performed several times daily in the ICU and used the GCS to assess patients’ conscious state. EEG data were recorded continuously (Cz reference) at a sampling rate of 250 Hz for several days or longer while patients were in the ICU. After data acquisition with Persyst software (Persyst Development Corporation, Solana Beach, CA, USA), data were exported in EDF format to MATLAB (The MathWorks, Inc., Natick, MA, USA) for analysis.

To analyze patients during periods of both high responsiveness (conscious) and minimal responsiveness (unconscious), we extracted 60 minutes of EEG from 13 channels common to all patients (Fp1, Fp2, F7, F8, T3, C3, Cz, C4, T4, T5, O1, O2, T6) at timepoints corresponding to consciousness, defined as GCS motor score ≥ 5 or GCS verbal score ≥ 4 (46, 47), and unconsciousness. EEG sections were spaced a minimum of 12 hours apart according to the following procedure, applied separately for conscious and unconscious data: 1) sorting each patient’s GCS scores from or high to low (conscious) or low to high (unconscious), 2) appending the highest (conscious) or lowest (unconscious) score to a second list, and 3) crawling down the first list of GCS scores and adding each timepoint that was at least 12 hours from any timepoint on the second list to the second list. 60-minute EEG sections were then extracted from the second list’s timepoints in order to sample the desired periods of consciousness and unconsciousness. Data were split into 10-second trials, demeaned, detrended, and re-referenced to the common average. Data were then visually inspected for artifacts. Data from electrodes with consistent drift or motion artifacts were removed, and 10-second trials with large motion artifacts spanning multiple electrodes were removed. We then ran an independent components analysis on the data to remove ocular or cardiac artifacts.

## ACKNOWLEDGMENTS

This work was supported by the National Institutes of Health grant RO1 MH111737 awarded to M.D., the National Institute of Neurological Disorders and Stroke grant NS21135 awarded to R.T.K, and a Tiny Blue Dot Foundation grant awarded to M.M.M. We would like to thank Michael A. Silver for his feedback on this work, Igor V. Ovchinnikov for guidance on the physics of edge-of-chaos criticality, and Julie Ashworth for extensive technical help. We would additionally like to thank Norman Spivak for organizing the medication records of our subjects with disorders of consciousness, and Courtney Real, Vikesh Shrestha, and Jesus E. Ruiz Tejeda for helping with clinical data collection and curation.

## Notes

https://figshare.com/articles/software/Consciousness_is_supported_by_near-critical_cortical_electrodynamics/12949355

### Competing Interest Statement

The authors have declared no competing interest.

