## Supplementary Information for "Consciousness is supported by near-critical cortical electrodynamics"

### SUPPLEMENTARY FIGURES

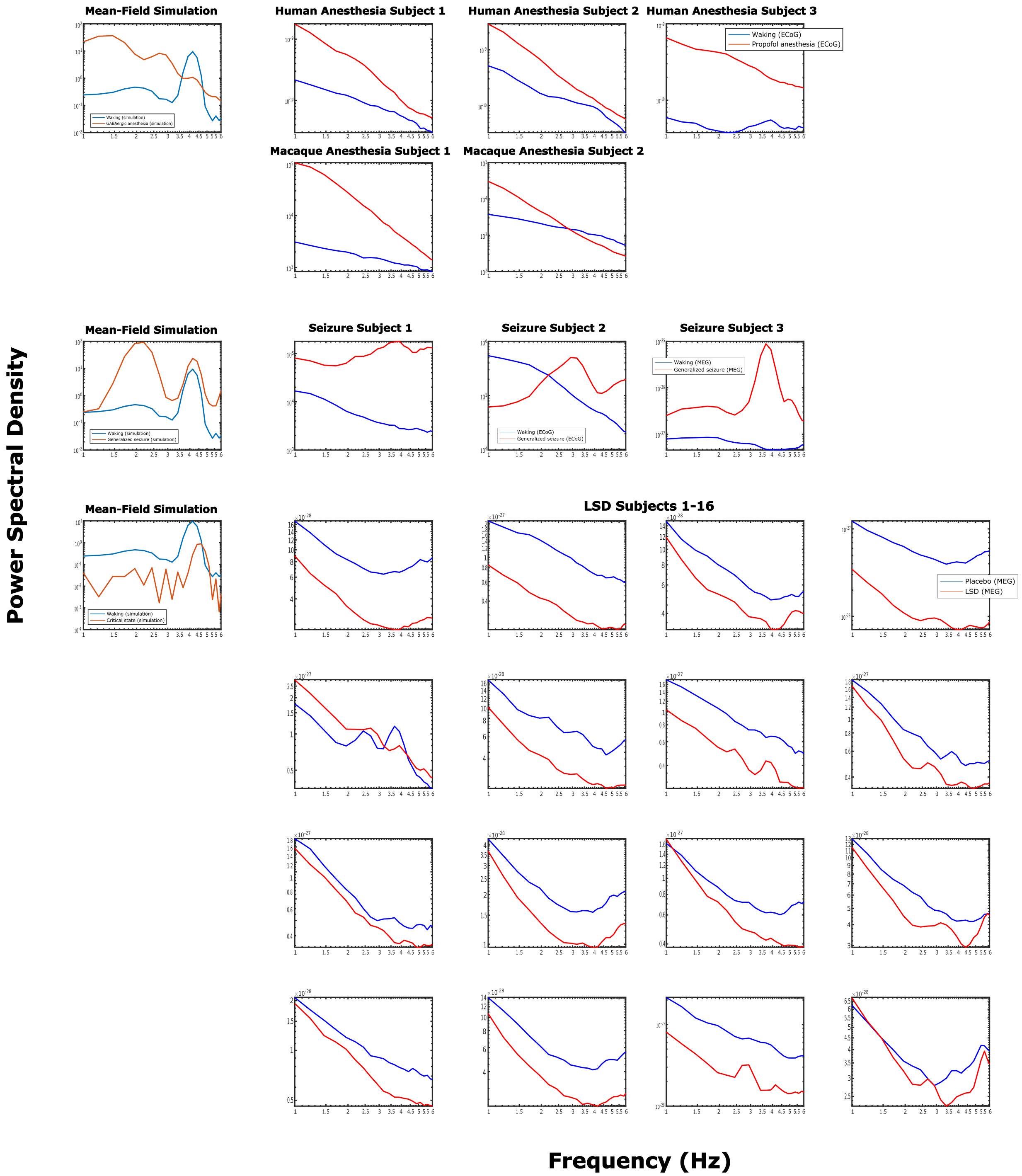


Fig. S1: (Caption on the following page.)

Fig. S1: We here plot the low-frequency power spectra of our real and simulated cortical electrodynamics in different states. For simulated data, we plot the median power spectrum across simulated cortical populations, and for empirical data, we plot the cross-channel median of each channel's cross-trial median power spectrum. Note that we here focus on 1-6 Hz, because the mean-field model only simulates oscillations intrinsic to the cortex in this range (without, for example, incorporating thalamic inputs, which are thought to underlie oscillations in the 8-12 Hz range). This was also the range over which we used the FOOOF algorithm to search for the frequencies of slow oscillations in our data. Note that the use of different datasets and imaging modalities led to wide variation in spectral power (for e.g., note that the spectral power of the MEG datasets was far lower than the spectral power of the ECoG datasets). In both the mean-field model and ECoG data, we see an increase in power and a steepening of spectral slope in the anesthesia state. Note that the model appears to have a peak at 4 Hz in the waking state that is absent in the real data, but that this apparent absence in real data is likely an artifact induced by averaging over channels whose slow oscillations extend over a range of frequencies (Figs. S2-S3). For the generalized seizure state, we see clear peaks between 3-4 Hz for both the model and real data. Though the model spectrum also includes a peak at 2 Hz that is apparently absent in the empirical data, note that this absence may again be an artifact of averaging over diverse channels (Figs. S2-S3). Finally, we compared power spectra in the LSD state to the power spectrum of the most information-rich and nearest-to-criticality state of the model (i.e., the only model parameter configuration that produced dynamics with a normalized Lempel-Ziv complexity greater than 0.99 for all three variants of Lempel-Ziv complexity as well as an absolute largest Lyapunov exponent less than 0.1). Like the LSD state, this parameter configuration - which corresponded to an increase in cortical excitability coupled with a slight decrease in the strength of gap junction coupling between inhibitory interneurons - resulted in reduced low-frequency power relative to the waking state. The similarities between the critical, maximally information-rich state of the model and the LSD state suggest possible follow-up work regarding potential molecular mechanisms of psychedelics - see Supplementary Note 2.

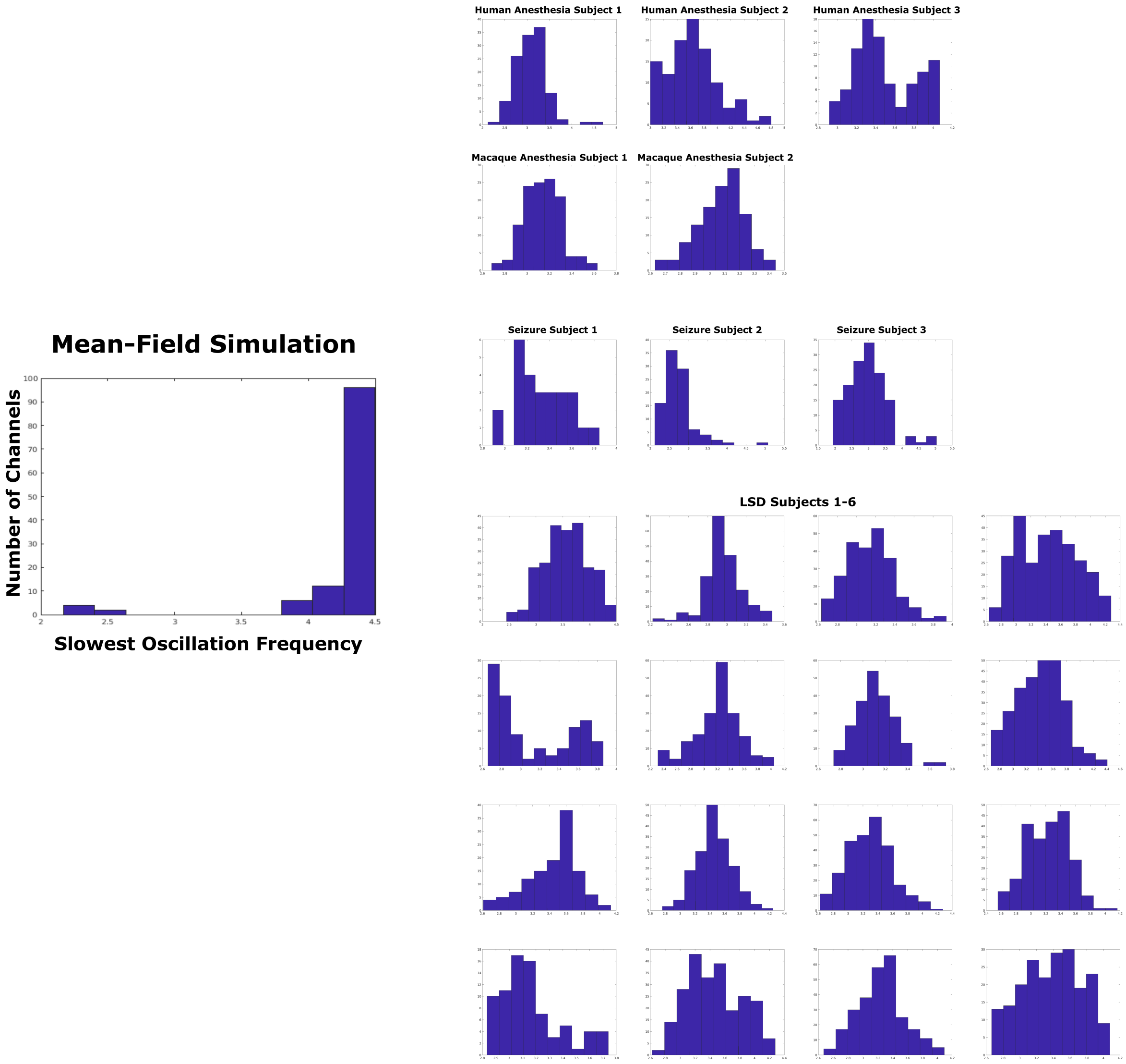


Fig. S2: (Caption on the following page.)

Fig. S2: The mean-field model used in this paper simulates the electrodynamics of a homogeneous patch of cortex, and as such produces electrodynamic oscillations with similar frequencies. Reflecting this, the FOOOF algorithm (as described in our Materials and Methods) identifies a relatively narrow range of slowest oscillation frequencies in the waking state of the model between 1-6 Hz (left). Our empirical recordings of cortical electrodynamics, however, were collected across heterogenous parts of cortex, and it is known that different parts of cortex produce electrodynamic oscillations at varying frequencies, which is one of the motivations behind the development of the FOOOF algorithm[1]. Thus, as expected, the FOOOF algorithm identified a broader range of slowest oscillation frequencies during normal waking states across the cortex of each one of our subjects (right). Therefore, in Fig. S3, we repeated the cross-state comparisons of power spectra as reported in Fig. S1, but looking exclusively at channels whose slowest oscillation frequencies during normal waking states were close to those of the waking state of the mean-field model.

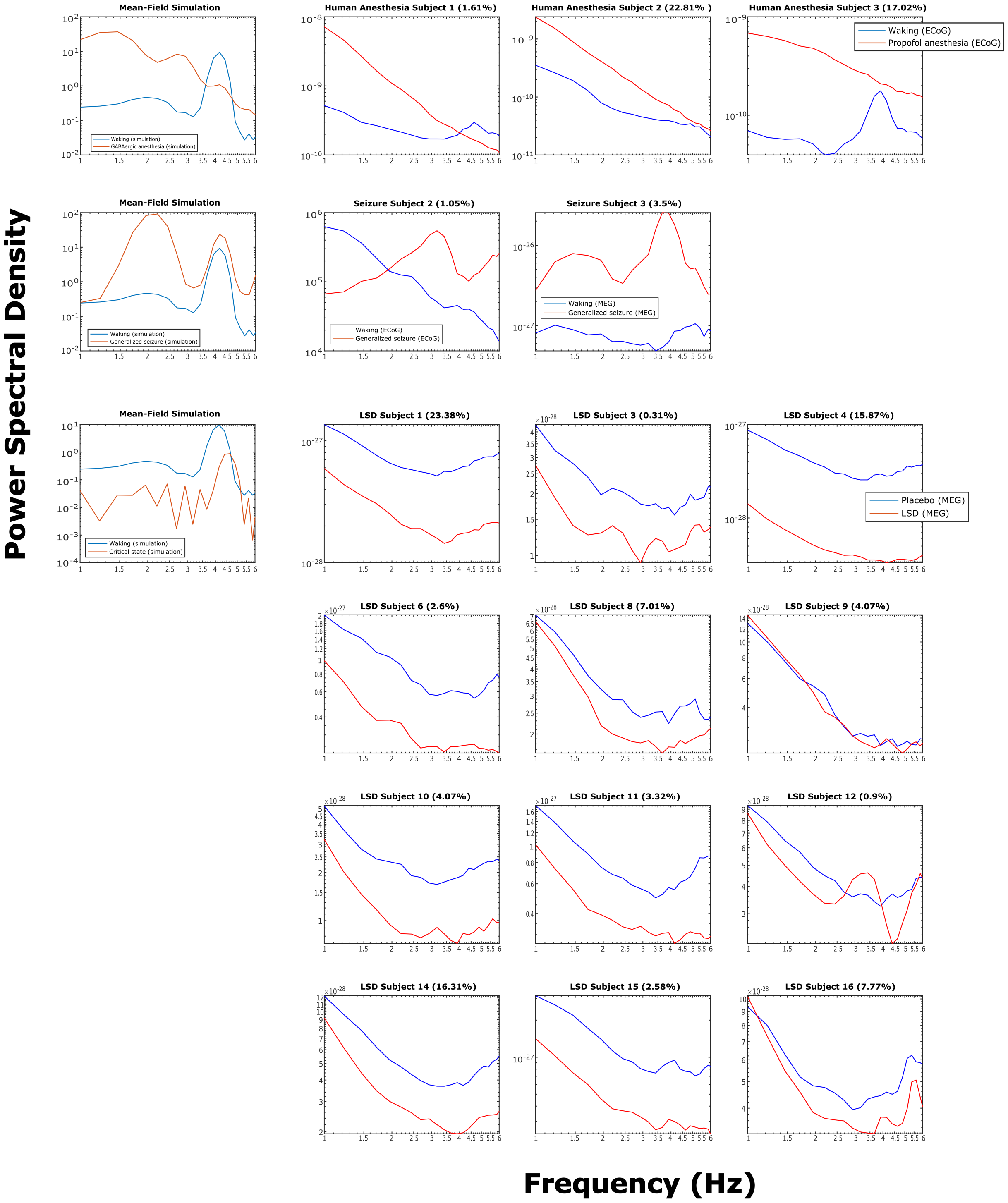


Fig. S3: (Caption on the following page.)

Fig. S3: To compare the power spectra of channels whose oscillatory behavior during normal waking states more closely matched that of the waking state of the mean-field model, we identified channels whose cross-trial median slowest oscillation frequencies during wake states (Fig. S3, right) were within $\pm 10\%$ of the median of the slowest oscillation frequencies of the waking state of the model (Fig. S2, left), which was 4.3 Hz. Note that several subjects had no channels whose slowest identified oscillation frequencies were within this range, and so are not plotted here. For each subject whose results are plotted here, we additionally indicate the percentage of their cortical channels that fell within this frequency range during normal waking states. Looking at just these channels, we found a 3-4 Hz rise above the background power spectrum during waking states for most subjects, which better matches the mean-field model, and also observed spectral changes during altered states (i.e., anesthesia, generalized seizures, and LSD) that likewise matched both the model and what we observed for the median across all channels (Fig. S1). We also found that the power spectrum of Seizure Subject 3's generalized seizure more closely matched the power spectrum of the simulated seizure, with peaks at both 1.5 Hz and 4 Hz.

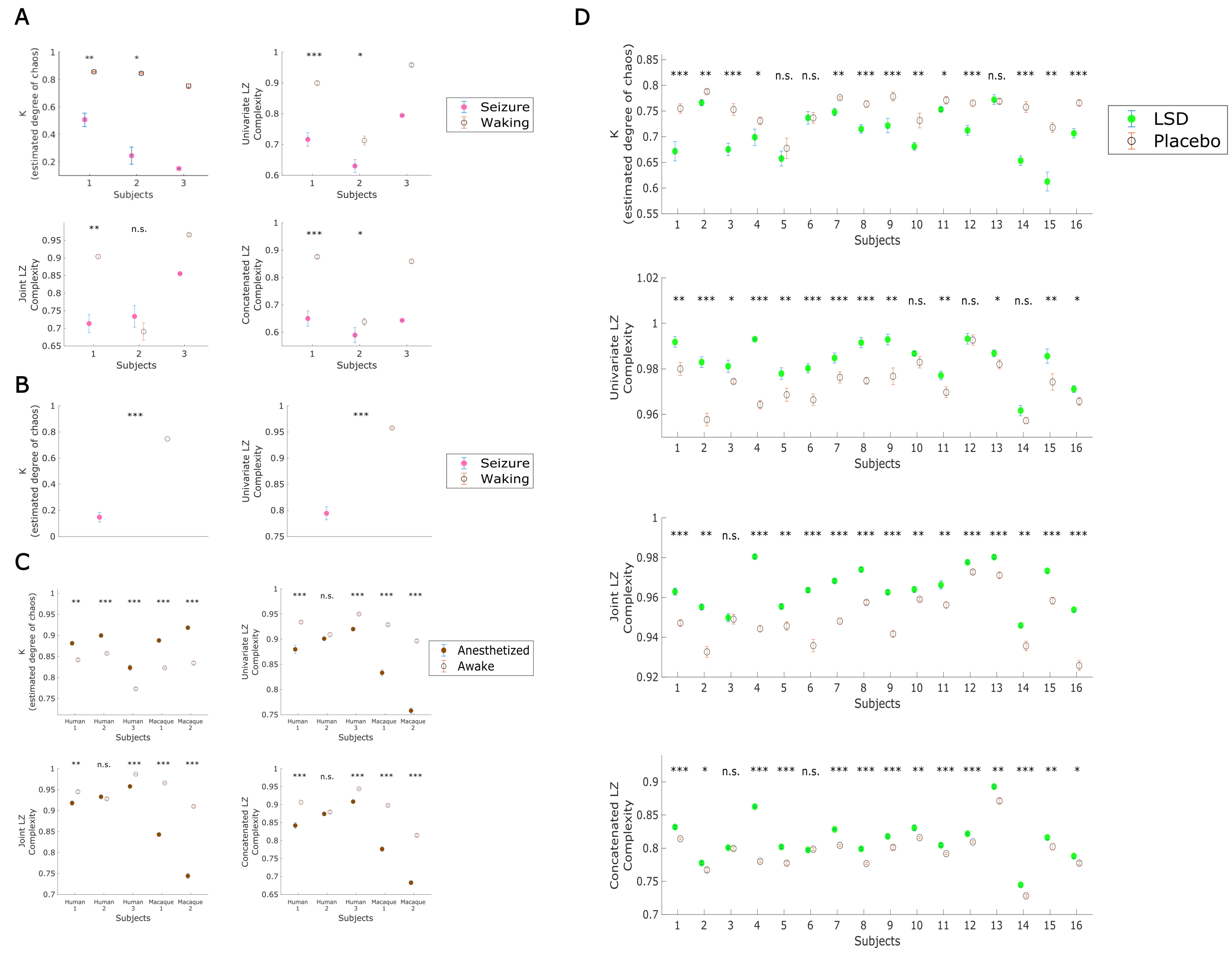


Fig. S4: (Caption on the following page.)

Fig. S4: **A**. We applied the modified 0-1 chaos test and three variants of Lempel-Ziv complexity to surface ECoG recordings from two human epilepsy patients (subjects 1 2) and to magnetoencephalography (MEG) recordings from another human epilepsy patient (subject 3) experiencing generalized seizures. Circles indicate the median estimated values across all trials per condition for a single subject, and errorbars indicate standard error of the median (estimated by taking the standard deviation of a bootstrap distribution of sample medians). Differences between conditions were tested using a left-tailed overlapping block bootstrap test (which controls for the non-independence of successive datapoints by preserving local time-series autocorrelations) with a block size of three trials (30 seconds of recording), to test against the null hypothesis that there is no decrease in the median K-statistic or Lempel-Ziv complexity during generalized seizures. The decreased chaoticity during generalized seizures was significant for both Subjects 1 and 2. Note that Subject 3 had only one $\sim$10 second generalized seizure during the duration of the recording, and so statistical comparison across trials was not possible - see **B** for statistical analysis of Subject 3's seizure on the level of individual MEG channels, rather than trials. Median information-richness, as measured by three different variants of Lempel-Ziv complexity, dropped significantly during generalized seizures for Seizure Subjects 1 and 2, with the exception of joint Lempel-Ziv complexity for Subject 2. Note again that no statistical analysis was performed for Seizure Subject 3 because the subject had only one generalized seizure; see **B** for analysis of the univariate Lempel-Ziv complexity of their individual channels across brain states. **B**. Here, we show the median and standard error of the median chaoticity (left) and univariate Lempel-Ziv complexity (right) across all channels during seizure subject 3's seizure trial and across their nine waking baseline trials. As was the case for the cross-trial comparisons (**A**), cross-channel differences between conditions were tested using left-tailed overlapping block bootstrap tests. The drop in the estimated chaoticity and information-richness of this subject's cortical electrodynamics was significant, consistent with the hypothesis that the dynamics of generalized seizures are information-poor and periodic/stable. **C**. We applied the modified 0-1 chaos test and the three variants of Lempel-Ziv complexity to surface ECoG recordings from three human subjects and two macaques under GABAergic anesthesia. Circles indicate the median values across all trials, per condition, and errorbars indicate standard error of the median. Differences in chaoticity between conditions were tested using a right-tailed overlapping block bootstrap test, to test against the null hypothesis of no increase in the median K-statistic under GABAergic anesthesia, and differences in information-richness were assessed using a left-tailed overlapping block bootstrap test, to specifically test against the null hypothesis that there is no decrease in Lempel-Ziv complexity during GABAergic anesthesia. The increased chaoticity under GABAergic anesthesia was significant for all subjects; the decrease in all measures of Lempel-Ziv complexity during anesthesia was significant for all subjects except Human Anesthesia Subject 2. **D**. We applied the modified 0-1 chaos test and Lempel-Ziv complexity algorithm to cortical MEG recordings from 16 human subjects following intravenous administration of either 75 $\mu$g of LSD or a saline placebo. Differences in chaoticity between conditions were tested using left-tailed overlapping block bootstrap tests, to test against the null hypothesis of no decrease in the median K-statistic during the psychedelic state relative to placebo, and differences in information-richness were assessed using a right-tailed overlapping block bootstrap test, to test against the null hypothesis of no increase in Lempel-Ziv complexity during the psychedelic state. The decreased chaoticity during the psychedelic state was significant for 13/16 subjects, univariate Lempel-Ziv complexity increased significantly for 15/16 subjects, joint Lempel-Ziv complexity increased significantly for 15/16 subjects, and concatenated Lempel-Ziv complexity increased significantly for 14/16 subjects. $*$ $p<0.05$, $**$ $p<0.01$, $***$ $p<0.001$.

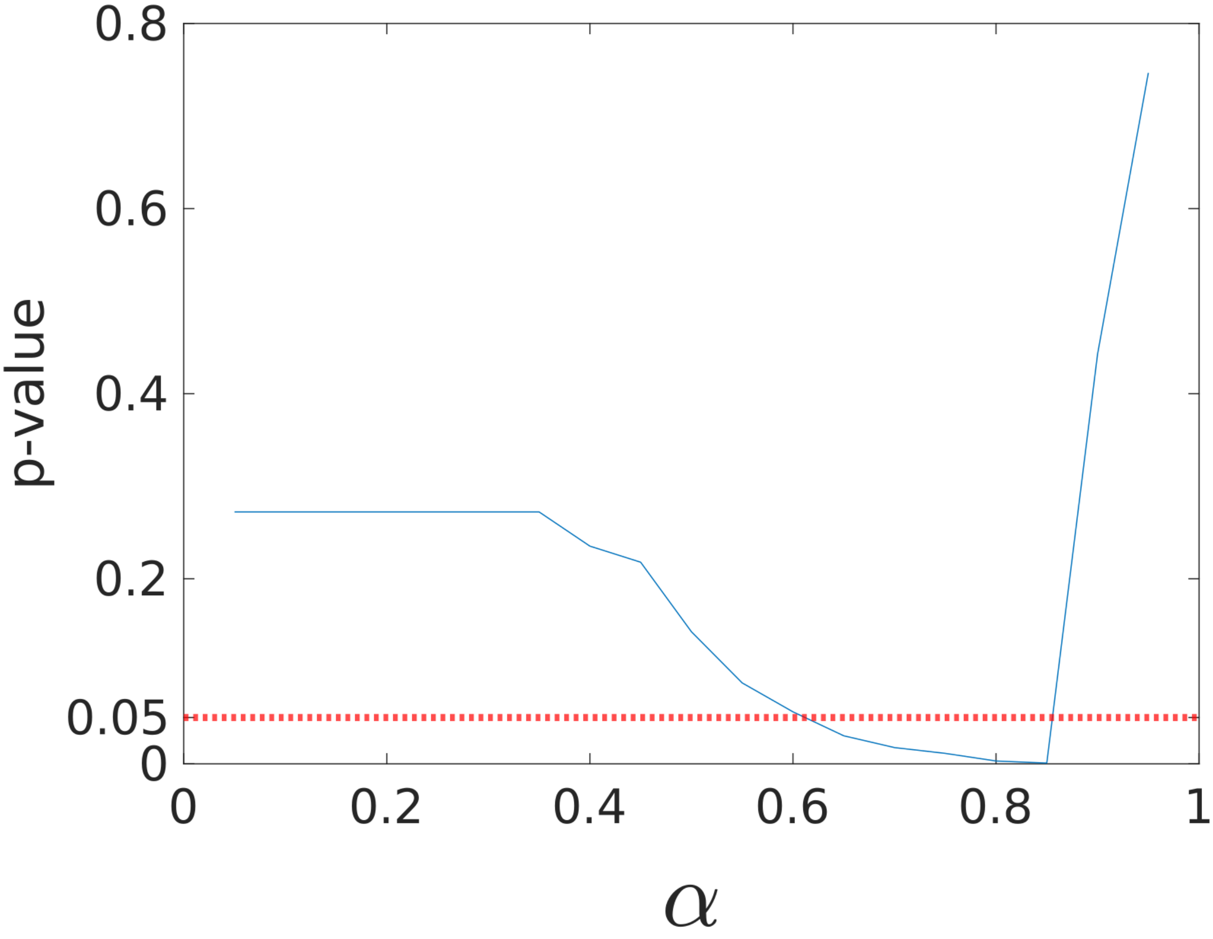


Fig. S5: Our new time-series estimate $c$ of proximity to edge-of-chaos criticality, based on a nonlinear transformation of the $K$-statistic outputted by the modified 0-1 chaos test, includes a parameter $\alpha$, which must be set between zero and one (see Materials and Methods). To explore the optimal range of $\alpha$ values for which $c$ might be used as a clinical biomarker of consciousness, we converted the cross-trial, state-specific medians of chaoticity estimates $K$ of 12 subjects for whom data were available from both conscious and unconscious states (namely five anesthesia subjects, three generalized seizure subjects, and four DOC patients) into our criticality measure $c$, using 19 unique values of $\alpha$. For each value of $\alpha$, we performed a right-tailed Wilcoxon rank sum test between the 12 values of $c$ from conscious states and the 12 values of $c$ from unconscious states, to test against the null hypothesis that there is no increase in median estimates of criticality during conscious states. We here plot the uncorrected p-values of these tests, for each value of $\alpha$ assessed. After conservative Bonferroni correction, the difference in $c$ at $\alpha$=0.85 between conscious and unconscious states remained significant, suggesting that $\alpha$ should be set around this value when exploring the potential utility of our new criticality measure $c$ as a biomarker of consciousness. We therefore set $\alpha$ to 0.85 in all subsequent analyses using our measure $c$ in both the main body of the paper and SI Appendix.

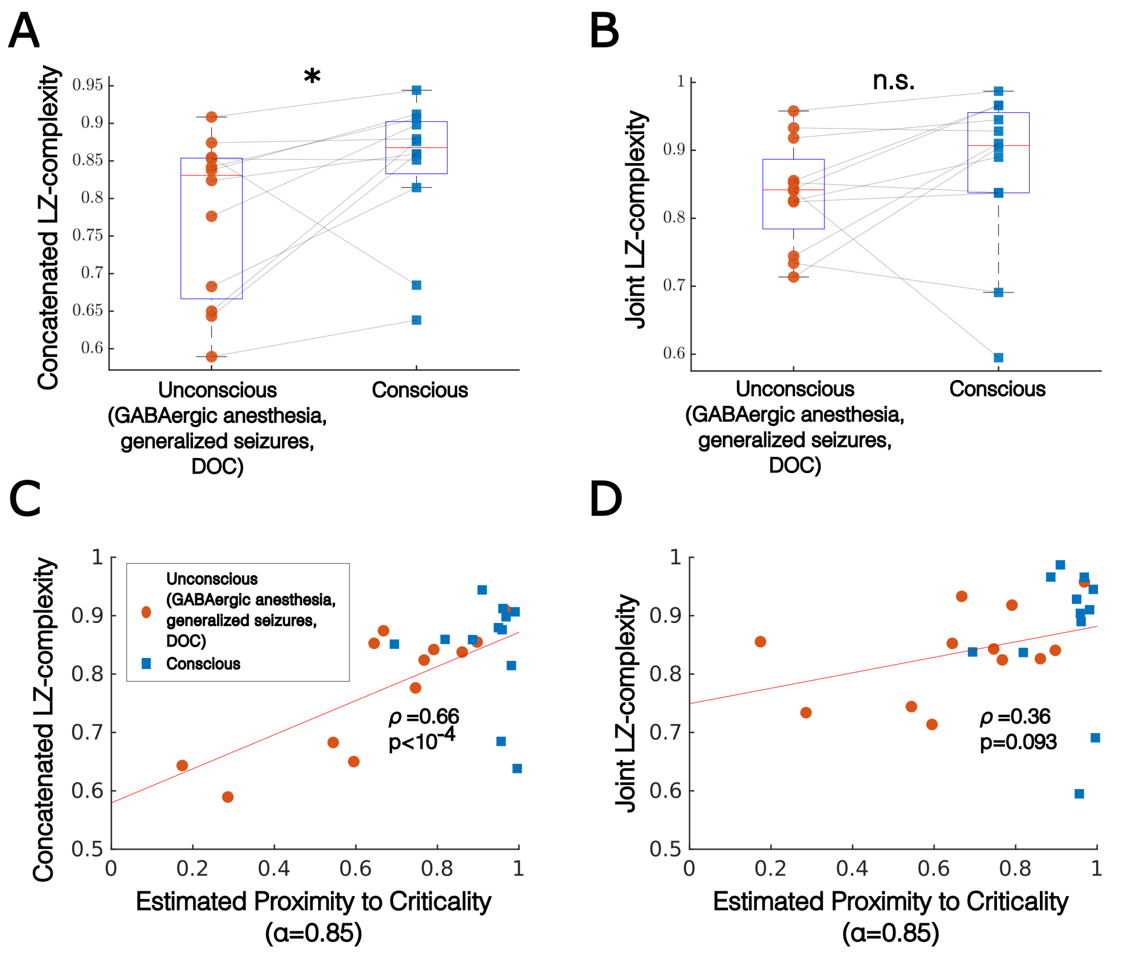


Fig. S6: **A** As was the case for univariate Lempel-Ziv complexity (Fig. 4B), concatenated Lempel-Ziv was significantly higher during conscious states than during unconscious states for the 12 subjects for whom data were available from both conscious and unconscious states (five anesthesia subjects, three generalized seizure subjects, and four DOC patients). Each dot represents the cross-trial median across a single subject's trials in either their unconscious or conscious state. Significance was assessed across subjects using a right-tailed Wilcoxon rank-sum test. **B** A cross-subject right-tailed Wilcoxon rank-sum test did not reveal a significant difference in joint Lempel-Ziv complexity across conscious vs. unconscious states. **C** As was the case for univariate Lempel-Ziv complexity (Fig. 4C), concatenated Lempel-Ziv complexity was significantly correlated with our criticality measure $c$ at $\alpha$=0.85 across subjects and states (partial correlation $\rho$=0.66, p$<10^{-4}$, controlling for median frequency at which signals were low-pass filtered). **D** Joint Lempel-Ziv complexity was not significantly correlated with our criticality measure.

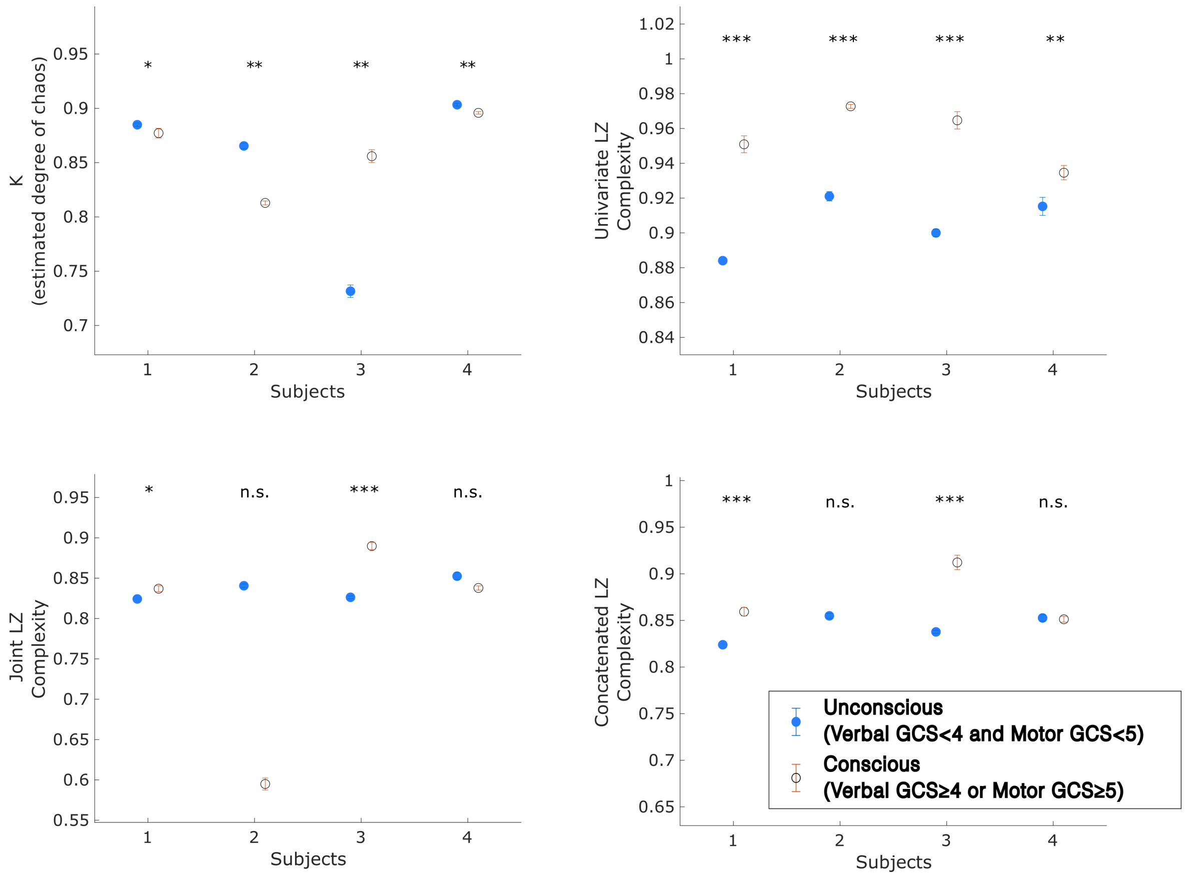


Fig. S7: We applied the modified 0-1 chaos test and the three variants of Lempel-Ziv complexity to clinical EEG recordings of the cortical electrodynamics of four human patients who recovered consciousness after going into a coma, following a traumatic brain injury. Circles indicate the median estimated chaoticity across all trials, per condition, and errorbars indicate standard error of the median. Differences in estimated chaoticity between conditions were tested using a two-tailed overlapping block bootstrap test (the test was two-tailed, rather than one-tailed, because we had no *a priori* prediction as to whether chaoticity would increase or decrease during unconscious relative to conscious states). Differences in information-richness were assessed using a left-tailed overlapping block bootstrap test, to specifically test against the null hypothesis that there is no decrease in Lempel-Ziv complexity in DOC. We found that estimated chaoticity (top left) was significantly higher in unconscious than conscious states for three out of four of the patients, similar to what we observed for GABAergic anesthesia (SI Appendix, Fig. S2C), but was significantly lower in the fourth patient, similar to what we observed for generalized seizures (SI Appendix, Fig. S2A-B). Univariate Lempel-Ziv complexity (top right) was significantly lower during unconscious than conscious states for all patients, recapitulating what we observed for both anesthesia and generalized seizures (SI Appendix, Fig. S2), while results with both measures of multivariate Lempel-Ziv complexity were inconsistent across patients (bottom).

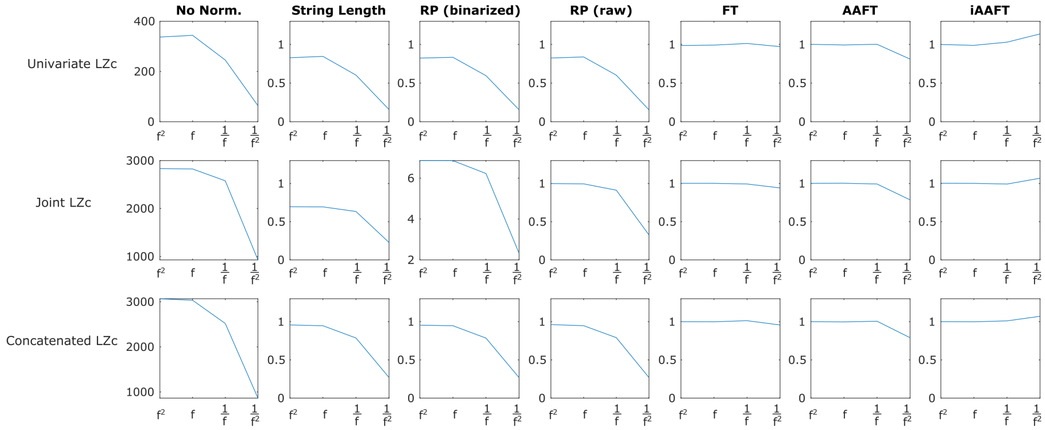


Fig. S8: In order to compare the amount of non-redundant information in different signals, Lempel-Ziv complexity is typically normalized to lie roughly between 0 and 1. We here test several different normalization approaches by applying them to noise signals with different spectral slopes; this is important in comparing Lempel-Ziv complexity across different brain states, as the spectral profiles of brain signals can vary significantly across different brain states. The x-axes here refer to multivariate noise signals with increasingly negative spectral slopes: $f^2$ (violet) noise, $f$ (blue) noise, $1/f$ (pink) noise, and $1/f^2$ (red) noise. All of these signals are entirely random (and thus contain entirely non-redundant information), and vary only by their spectral properties; thus, a properly normalized measure of Lempel-Ziv complexity should be constant (and approaching the ceiling of 1) for each of these random processes. The first column ("No Norm.") shows the behavior of our three measures of Lempel-Ziv complexity as a function of the spectral profile of the noise signals; note the drop as the power spectrum steepens from pink ($1/f$) noise to red ($1/f^2$) noise. The second column, "String Length," shows the behavior of the three complexity measures normalized by $T/\log_2 T$, where $T$ is the length of the binary string being compressed; note that this form of normalized Lempel-Ziv complexity likewise varies as a function of the spectral slope of the random signals. The third column, "RP (binarized)," shows the alternative normalization scheme used by Schartner and colleagues[2,3], which divides the Lempel-Ziv complexity of a signal (after it has been binarized) by the Lempel-Ziv complexity of a random shuffle of the binarized form of the signal; this likewise is affected by spectral slope. The fourth column, "RP (raw)" is similar, but instead divides the Lempel-Ziv complexity of a signal by the Lempel-Ziv complexity of a random shuffle of that continuous signal (i.e., Lempel-Ziv complexity is calculated for a binarized version of the original signal, and for a binarized version of the shuffled signal, and the former is normalized by the latter). The final three columns show the behavior of Lempel-Ziv complexity of a signal divided by the Lempel-Ziv complexity of a phase-randomized surrogate of that signal, following the methodology of Brito and colleagues[4]. A phase-randomized surrogate preserves the spectral properties of the original signal but is otherwise random. We used three phase-randomized surrogate algorithms: Fourier transform surrogates ("FT"), amplitude adjusted Fourier transform surrogates ("AAFT") and iterative amplitude adjusted Fourier transform surrogates ("iAAFT"). Normalizing by the Lempel-Ziv complexity of surrogates generated by any of these three algorithms produced desirable behavior - namely, a consistent normalization to near one for any noise signal, regardless of its spectral properties - but the least variance as a function of spectral slope was produced by Fourier transform surrogates ("FT"), and so we used this algorithm throughout the rest of this paper to create surrogates by which to normalize Lempel-Ziv complexity.

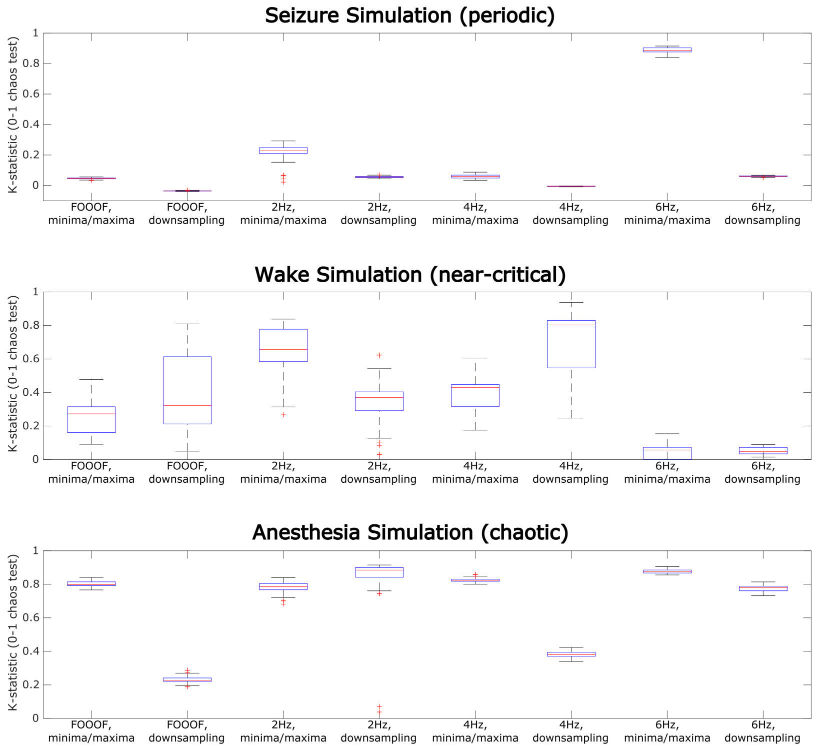


Fig. S9: To ensure that our processing pipeline produces estimates of chaoticity that are both accurate and stable, we ran 100 simulations of the mean-field model in its seizure (periodic), waking (near-critical), and anesthetized (chaotic) states. Each simulation included unique noise inputs. Thus, the deterministic component of all 100 simulations in a given state had the same ground-truth largest Lyapunov exponent (i.e. level of chaoticity), but the actual time-series produced in each simulation would be unique, due to the unique noise inputs. As in Table S2, we compared results using four low-pass filtering methods (namely, where the low-pass filter frequency was determined either by the FOOOF algorithm or set at 2, 4, or 6 Hz) and two different methods for time-discretization (namely, downsampling or taking the local minima and maxima) of the low-pass filtered signal. For any given low-pass filtering and time-discretization method, results using the modified 0-1 chaos test were highly stable in both the seizure (periodic) and anesthesia (chaotic) states. With the exception of highly stable results when low-pass filtering at 6 Hz (which, despite this stability, yielded inaccurate estimates of chaoticity, Table S2), results were somewhat less stable for the waking (near-critical) state, though we note the relative stability of the method used in this paper, based on simulation results (Table S2), i.e., taking the local minima and maxima of a time-series that was low-pass filtered at a frequency determined using the FOOOF algorithm. Given both the accuracy and relative stability of chaoticity estimates using this method, the chaoticity of all empirical time-series in this paper was estimated using the local minima and maxima of signals low-pass filtered using the FOOOF algorithm.

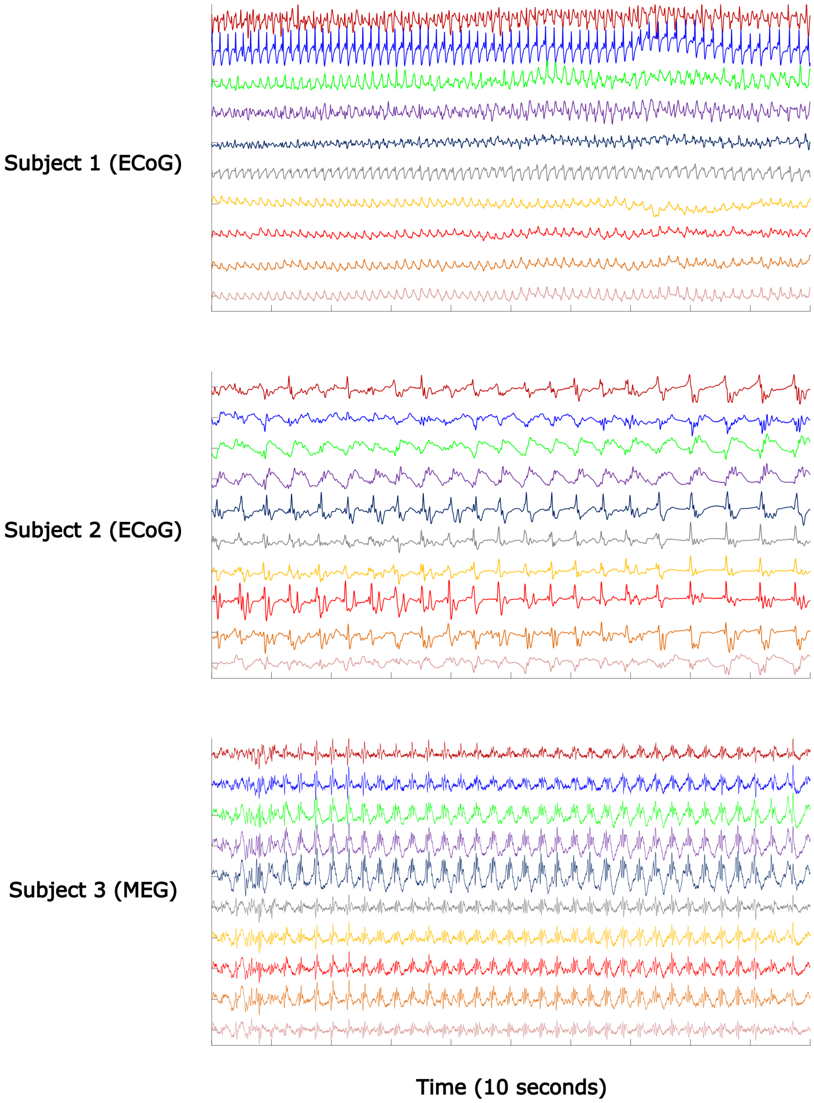


Fig. S10: 10-second time-traces from 10 sample channels during generalized seizures from our three seizure subjects. The subject numbers match those in figs. S1-S2.

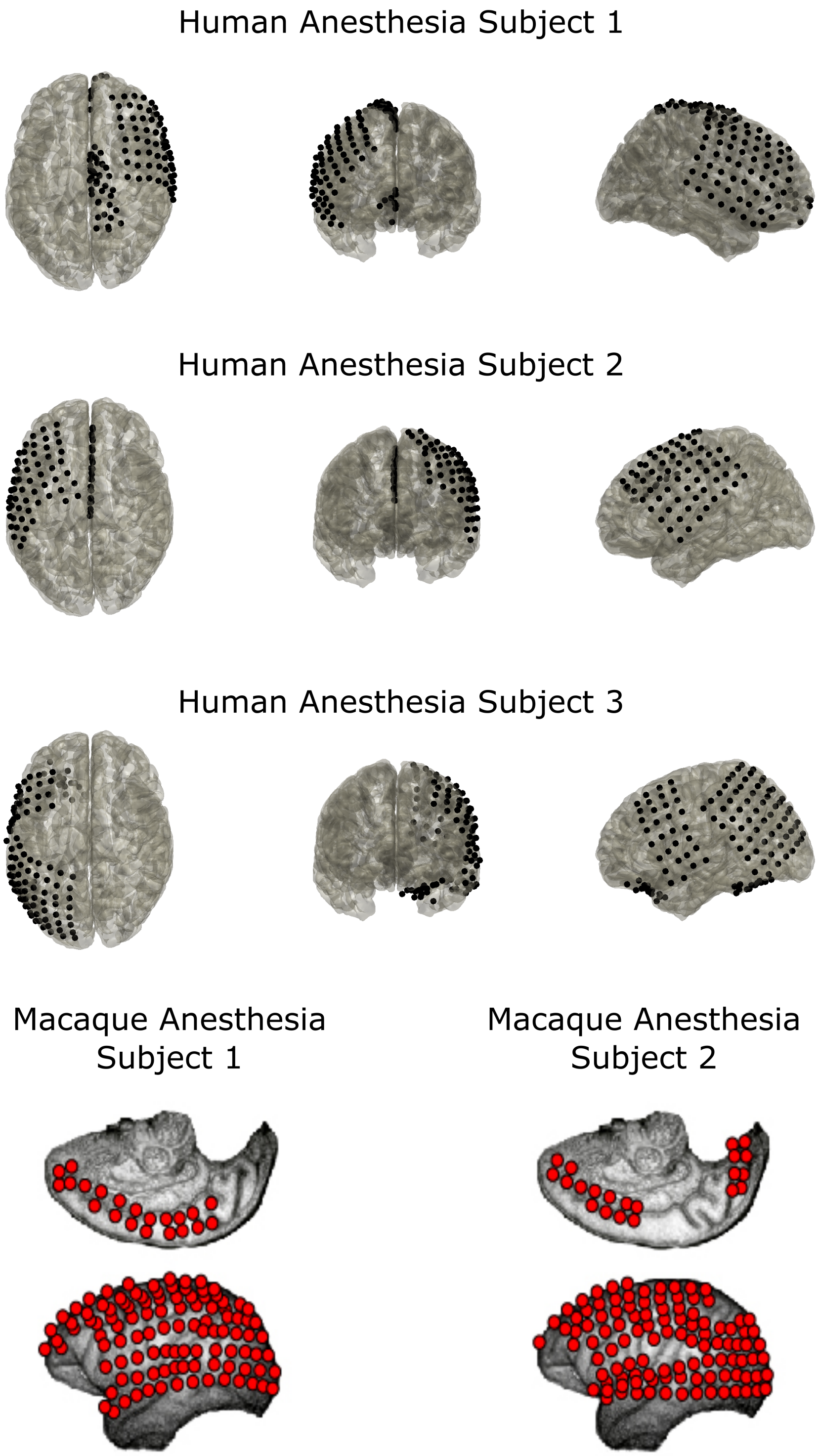


Fig. S11: Surface electrocorticography (ECoG) electrode placement for our anesthesia subjects.

### SUPPLEMENTARY TABLES

Table S1: We previously described[5] a test to determine whether a signal is generated by a predominantly deterministic system (linear or nonlinear). The test compares the permutation entropy of a signal to the permutation entropies of 1,000 amplitude adjusted Fourier transform surrogates and 1,000 cyclic phase permutation surrogates of that signal. If the permutation entropy of the original signal falls within either surrogate distribution, then the signal is classified as predominantly or "operationally" stochastic. We identified a failure case of this method, which is that for short, noise-free signals sampled at a high frequency from deterministic, continuous-time systems, the cyclic phase permutation algorithm generates surrogates with identical permutation entropies. This in turn can lead to misclassifications of deterministic processes as stochastic. We here modify our original test to catch this edge case: if the cyclic phase permutation algorithm fails to generate surrogates with at least 500 unique permutation entropy values, then jitter in the form of 2.5% white noise is incrementally added to the original signal until this failure case is broken. We here tested the accuracy of both our original and modified test on a broad range of dynamical systems, the code for which is included alongside our previously described Chaos Decision Tree Algorithm (see our prior work for model equations and parameters[5]). Note that we excluded one system described in our previous work, namely the noise-driven sine map, because it is unclear whether its dynamics are in fact predomoninantly stochastic, or whether they are better characterized as noise-induced tunneling between quasi-stable, deterministic attractor states[5]. We also applied both our original and modified stochasticity test to all 120 nodes of the mean-field model studied here, with its noise inputs turned off (so its dynamics are fully deterministic), in its seizure, waking, and anesthesia states. We found that our modified stochasticity test performed as well as the original test for all systems described in our previous work, and markedly outperformed the original test for the mean-field simulations of low-frequency cortical activity. We therefore applied this modified test to our real cortical recordings (Table S2). Table on next page.

| | Original Stochasticity Test Accuracy | Modified Stochasticity Test Accuracy |
|---|---|---|
| ARMA process (linear stochastic) | 67/100 (67%) | 71/100 (71%) |
| Bounded random walk (nonlinear stochastic) | 100/100 (100%) | 100/100 (100%) |
| Cubic Map (chaotic) | 100/100 (100%) | 100/100 (100%) |
| Cubic Map (Heagey-Hammel strange nonchaotic) | 100/100 (100%) | 100/100 (100%) |
| Cubic Map (periodic-doubled) | 100/100 (100%) | 100/100 (100%) |
| Cubic Map (periodic) | 100/100 (100%) | 100/100 (100%) |
| Cubic Map (type-3 intermittency strange nonchaotic) | 100/100 (100%) | 100/100 (100%) |
| Cyclostationary process (linear stochastic) | 98/100 (98%) | 98/100 (98%) |
| Freitas map (nonlinear stochastic) | 99/100 (99%) | 99/100 (99%) |
| Generalized Hénon Map (hyperchaotic) | 100/100 (100%) | 100/100 (100%) |
| GOPY map (strange nonchaotic) | 100/100 (100%) | 100/100 (100%) |
| Granulocyte levels (chaotic) | 100/100 (100%) | 100/100 (100%) |
| Granulocyte levels (periodic) | 100/100 (100%) | 100/100 (100%) |
| Hénon map (periodic) | 100/100 (100%) | 100/100 (100%) |
| Ikeda map (chaotic) | 100/100 (100%) | 100/100 (100%) |
| Logistic map (chaotic) | 100/100 (100%) | 100/100 (100%) |
| Logistic map (periodic) | 100/100 (100%) | 100/100 (100%) |
| Lorenz system (chaotic) | 100/100 (100%) | 100/100 (100%) |
| Poincaré oscillator (chaotic) | 100/100 (100%) | 100/100 (100%) |
| Poincaré oscillator (periodic) | 100/100 (100%) | 100/100 (100%) |
| Poincaré oscillator (quasi-periodic) | 100/100 (100%) | 100/100 (100%) |
| Random walk (linear stochastic) | 100/100 (100%) | 100/100 (100%) |
| Rössler oscillator (chaotic) | 100/100 (100%) | 100/100 (100%) |
| Simulated anesthesia state (chaotic) | 0/120 (0%) | 117/120 (98%) |
| Simulated seizure (periodic) | 0/120 (0%) | 120/120 (100%) |
| Simulated waking state (chaotic, near-critical) | 1/120 (1%) | 98/120 (82%) |
| Spiking Izhikevich neuron (chaotic) | 100/100 (100%) | 100/100 (100%) |
| Trended random walk (linear stochastic) | 100/100 (100%) | 99/100 (99%) |
| Unfiltered blue noise (linear stochastic) | 100/100 (100%) | 100/100 (100%) |
| Unfiltered pink noise (linear stochastic) | 100/100 (100%) | 100/100 (100%) |
| Unfiltered red noise (linear stochastic) | 100/100 (100%) | 100/100 (100%) |
| Unfiltered violet noise (linear stochastic) | 100/100 (100%) | 100/100 (100%) |
| Unfiltered white noise (linear stochastic) | 100/100 (100%) | 100/100 (100%) |

Table S2: To test whether low-frequency cortical electrodynamics are predominantly deterministic (which is an important assumption of the 0-1 chaos test), as well as to test against the possibility that our results were potentially driven by changing levels of stochasticity in low-frequency cortical dynamics rather than by changing levels of chaoticity, we applied the modified test of signal stochasticity described in Table S1 to all of our empirical datasets. Specifically, we applied the test to all data (channels x trials) low-pass filtered using the FOOOF algorithm (i.e., the data for which we assessed low-frequency chaoticity using the 0-1 test). We found that low-frequency dynamics from a majority (75%) of channels(x)trials were classified as predominantly deterministic. Moreover, a cross-subject, two-tailed Wilcoxon rank-sum test comparing percentage of channels(x)trials classified as stochastic in normal waking versus altered states revealed no significant difference between conditions (p=0.2446). Additionally, there was no correlation between the change in each subject's median K-statistic across conditions and the change in the percentage of their channels(x)trials that were classified as stochastic (Pearson correlation r=-0.304, p=0.116; partial correlation $\rho$=-0.189, p=0.346, controlling for change in median frequency at which signals were low-pass filtered). To further test against the possibility that these findings were potentially driven by our low-pass filtering method, we generated 100 simulations of 5,000 time-points of violet, blue, white, pink, and red noise, and applied this stochasticity test to these simulated noise samples after low-pass filtering in the 1-6 Hz range using the FOOOF algorithm (note that, as was the case for the empirical data, a simulated noise signal was excluded if the FOOOF algorithm did not identify an oscillation in this frequency range). Although our stochasticity test performed with perfect accuracy in classifying unfiltered noise as stochastic (Table S1), performance did drop for low-pass filtered noise. That said, we note that performance was very high for both filtered pink and red noise, whose power spectra ($\frac{1}{f}$ and $\frac{1}{f^2}$, respectively) more closely match that of real neural electrodynamics[1] than do those of other noise colors. Taken together, these results suggest that low-frequency cortical electrodynamics are predominantly deterministic, and that observed changes in the K-statistic across brain states result from changing levels of chaoticity in those electrodynamics, as predicted, rather than changing levels of stochasticity. Table on next page.

| Subject | Fraction of channels(x)trials classified as stochastic (low-frequency) | |
|---|---|---|
| | Normal waking | Altered state |
| Anesthesia Subject 1 (ECoG, human) | 3770/5421 (70%) | 5499/6522 (84%) |
| Anesthesia Subject 2 (ECoG, human) | 8538/11178 (76%) | 9121/10853 (84%) |
| Anesthesia Subject 3 (ECoG, human) | 5070/10041 (50%) | 4423/8030 (55%) |
| Anesthesia Subject 4 (ECoG, macaque) | 264/11948 (2%) | 0/13528 (0%) |
| Anesthesia Subject 5 (ECoG, macaque) | 226/11772 (2%) | 4/10249 (0%) |
| Seizure Subject 1 (ECoG, human) | 9/1539 (1%) | 0/225 (0%) |
| Seizure Subject 2 (ECoG, human) | 0/1385 (0%) | 0/1128 (0%) |
| Seizure Subject 3 (MEG, human) | 208/1007 (21%) | 54/143 (38%) |
| LSD Subject 1 (MEG, human) | 2245/8677 (26%) | 1445/9870 (15%) |
| LSD Subject 2 (MEG, human) | 2689/10081 (27%) | 768/6548 (12%) |
| LSD Subject 3 (MEG, human) | 2616/12679 (21%) | 2128/12017 (18%) |
| LSD Subject 4 (MEG, human) | 2674/13300 (20%) | 1036/13732 (8%) |
| LSD Subject 5 (MEG, human) | 1644/6265 (26%) | 2226/6343 (35%) |
| LSD Subject 6 (MEG, human) | 1854/7448 (25%) | 1230/10564 (12%) |
| LSD Subject 7 (MEG, human) | 2532/10526 (24%) | 1843/10549 (17%) |
| LSD Subject 8 (MEG, human) | 2658/10412 (26%) | 2236/9667 (23%) |
| LSD Subject 9 (MEG, human) | 795/3193 (25%) | 558/2824 (20%) |
| LSD Subject 10 (MEG, human) | 486/6344 (8%) | 376/5946 (6%) |
| LSD Subject 11 (MEG, human) | 3118/12201 (26%) | 3059/13317 (23%) |
| LSD Subject 12 (MEG, human) | 1545/8818 (18%) | 1406/8866 (16%) |
| LSD Subject 13 (MEG, human) | 63/1018 (6%) | 54/939 (6%) |
| LSD Subject 14 (MEG, human) | 1964/8765 (22%) | 1432/7688 (19%) |
| LSD Subject 15 (MEG, human) | 2806/9350 (30%) | 1438/10028 (14%) |
| LSD Subject 16 (MEG, human) | 1310/7855 (17%) | 1132/6439 (18%) |
| Coma patient 1 (EEG, human) | 360/3230 (11%) | 1481/11914 (12%) |
| Coma patient 2 (EEG, human) | 6187/12138 (51%) | 1643/3904 (42%) |
| Coma patient 3 (EEG, human) | 221/416 (53%) | 12538/28752 (44%) |
| Coma patient 4 (EEG, human) | 1174/2565 (46%) | 869/2199 (40%) |
| Filtered violet noise | 82/83 (99%) | |
| Filtered blue noise | 61/94 (65%) | |
| Filtered white noise | 33/88 (38%) | |
| Filtered pink noise | 65/93 (70%) | |
| Filtered red noise | 91/92 (99%) | |

Table S3: We here compare different time-series analysis methods for tracking changing levels of chaoticity in a system. To do so, we assessed the Pearson correlation $r$ and associated p-values (Bonferroni-corrected) between the ground-truth largest Lyapunov exponent of the mean-field cortical model studied in this paper (calculated with the noise input turned off) and the K-statistic of the 0-1 chaos test, applied to the time-discretized, low-frequency component of the simulated dynamics of the model (with the noise input turned on, so as to better assess their ability to track changing chaoticity in real cortical dynamics). Low-frequency dynamics were extracted using EEGLAB's two-way least-squares FIR low-pass filtering, as described in the Materials and Methods. Cutoff frequencies for low-pass filtering were determined in a data-driven manner using the FOOOF algorithm, or were set at 2 Hz, 4 Hz, or 6 Hz. Once data were low-pass filtered, they were time-discretized using either the iterative downsampling method described in our prior work[5], or by taking all local minima and maxima of the simulated signals[6]. We further tested the ability of these analysis methods to track varying degrees of chaos in the presence of very high levels of either white or pink (1/f) measurement noise, where the noise amplitude equaled half (50%) the standard deviation of the underlying signal. Overall, we found that the K-statistic, when applied to low-frequency activity extracted using the FOOOF algorithm and discretized using the local minima/maxima method (bolded), outperformed all other methods in tracking the ground-truth chaoticity of the model, and was unaffected by even large amounts of measurement noise. This method also produced stable estimates of chaoticity across unique model simulations (Fig. S7). We therefore used this method to track changing levels of chaoticity in our real cortical recordings.

| | Noise-free | | 50% White Noise | | 50% Pink (1/f) Noise | |
|---|---|---|---|---|---|---|
| | r | p-value | r | p-value | r | p-value |
| **FOOOF, minima/maxima** | **0.84** | **<10e-4** | **0.84** | **<10e-4** | **0.83** | **<10e-4** |
| FOOOF, downsampling | 0.72 | <10e-4 | 0.72 | <10e-4 | 0.73 | <10e-4 |
| 2 Hz, minima/maxima | 0.75 | <10e-4 | 0.72 | <10e-4 | 0.38 | <10e-4 |
| 2 Hz, downsampling | 0.16 | <10e-4 | 0.1 | n.s. | 0.06 | n.s. |
| 4 Hz, minima/maxima | 0.74 | <10e-4 | 0.74 | <10e-4 | 0.75 | <10e-4 |
| 4 Hz, downsampling | 0.73 | <10e-4 | 0.73 | <10e-4 | 0.67 | <10e-4 |
| 6 Hz, minima/maxima | -0.20 | <10e-4 | -0.22 | <10e-4 | -0.39 | <10e-4 |
| 6 Hz, downsampling | 0.73 | <10e-4 | 0.72 | <10e-4 | 0.47 | <10e-4 |

Table S4: Because the FOOOF algorithm picks a unique frequency at which to low-pass filter data for each channel and trial, we here calculated the partial correlation $\rho$ (and Bonferroni-corrected p-values) between the ground-truth largest Lyapunov exponents of the mean-field model and the K-statistic calculated using the FOOOF algorithm as described above, controlling for the low-pass filter frequency selected by the FOOOF algorithm. We found that even when controlling for the frequency selected by the FOOOF algorithm for low-pass filtering, the method identified in Table S2 as superior to all alternatives - namely, applying the 0-1 chaos test to the local minima and maxima of a signal low-pass filtered at a frequency identified by the FOOOF algorithm - robustly tracks ground-truth chaoticity. We moreover note that, unlike the minima/maxima time-discretization method, controlling for frequency markedly reduced the correlation between ground-truth largest Lyapunov exponents and the K-statistic calculated using the iterative down-sampling method.

| | Noise-free | | 50% White Noise | | 50% Pink (1/f) Noise | |
|---|---|---|---|---|---|---|
| | $\rho$ | p-value | $\rho$ | p-value | $\rho$ | p-value |
| **FOOOF, minima/maxima** | **0.82** | **$<$10e-4** | **0.81** | **$<$10e-4** | **0.82** | **$<$10e-4** |
| FOOOF, downsampling | 0.56 | $<$10e-4 | 0.57 | $<$10e-4 | 0.59 | $<$10e-4 |

Table S5: All four DOC patients whose EEG recordings were analyzed in this paper received painkillers and anesthetics as part of their treatment protocol. Unfortunately, it was not possible to determine the exact timing of drug administration relative to behavioral assessment using the Glasgow Coma Scale (GCS) and corresponding EEG recordings, and so our results for these patients could be confounded by the effects of these drugs. Here, we list the drugs that were administered to each patient *on the same day as* GCS scoring and corresponding EEG recordings, which are the only data that are available regarding the relative timing of drug administration and GCS assessment.

| **Subjects** | **Unconscious** | **Conscious** |
|---|---|---|
| 1 | Propofol, ketamine, dexmedetomidine | none |
| 2 | Propofol, opioids, ketamine, dexmedetomidine, benzodiazepines | none |
| 3 | Propofol, opioids, ketamine, dexmedetomidine, barbituates, benzodiazepines | Propofol, opioids, ketamine, dexmedetomidine, benzodiazepines |
| 4 | Propofol, opioids, ketamine, dexmedetomidine, barbituates | Propofol, opioids, ketamine, dexmedetomidine, barbituates |

### SUPPLEMENTARY NOTES

**Supplementary Note 1**

While “chaos” and “disorder” have often been used interchangeably in the literature on neural criticality, this is in fact misleading: counter-intuitively, chaos is in fact the “ordered” phase of a dynamical system with respect to edge-of-chaos criticality[7,8]. This is because a phase is considered “ordered” with respect to a critical point if it is characterized by whatever symmetry is broken at that critical point, and it is known that the topological or de-Rahm supersymmetry of dynamical systems is broken at the periodic-to-chaotic transition[7–14]. A more intuitive example of this is ice water, which is “ordered” because it lacks the translational and rotational symmetry that makes liquid water “disordered.” In other words, because liquid water is disordered, any rotation or translation (along any axis) would preserve the shape of the water, meaning that it has both translational and rotational symmetry; in contrast, because ice water has a regular crystalline lattice structure, that structure will only be preserved for a limited number of rotations or translations, which means that ice water lacks translational and rotational symmetry. This lack of symmetry is what makes ice water “ordered.” Similarly, because periodic systems are topologically symmetric and because this symmetry is broken at the edge-of-chaos critical point, chaos is in fact the “ordered” phase of dynamical systems. Moreover, it is important to note that this topological or de-Rahm supersymmetry can be broken either by the non-integrability of a dynamical system (which is the classic definition of chaos in the deterministic sense) *or* by noise-induced tunneling between attractors (which extends the notion of chaos to stochastic dynamical systems)[7–14]. Thus, our finding that low-frequency cortical electrodynamics are weakly chaotic during conscious states is both fully consistent (if counter-intuitively so) with the proposal that waking cortical dynamics operate on the ordered side of criticality, and with the presence of dynamic noise in cortical networks.

**Supplementary Note 2**

We suggest a follow-up experiment to a result reported here, which is that the mean-field model of cortical electrodynamics exhibited its maximally information-rich, nearest-to-criticality behavior when there was a moderate reduction in the strength of gap junction coupling between inhibitory interneurons, as well as an increase in postsynaptic excitability - a parameter change which led to reduced spectral power at low frequencies (Fig. S1, S3), recapitulating results observed for the MEG recordings from humans following administration of LSD (Fig. S1, S3). This potentially points to an as-yet-unstudied molecular effect of psychedelics. It is already known that psychedelics induce a marked increase in the frequency and amplitude of spontaneous glutamatergic excitatory postsynaptic potentials in cortical layer V pyramidal cells[15–18], which leads to reduced low-frequency local field potential[19] and MEG power[20], and that an increase in cortical excitability in mean-field models can recreate many of the effects of psychedelics on macro-scale cortical dynamics[20,21]. But, there is currently no published research on the effect of psychedelics on cortical gap junction coupling[22]. That said, in light of both our simulation-based results and prior empirical findings, we predict an inhibitory effect of psychedelics on gap junction coupling. To begin, it is already known that serotonin and other 5-HT2A receptor agonists suppress gap junction coupling, and that 5-HT2A receptor antagonists attenuate this effect[23]. Additionally, the antipsychotic drugs clozapine and haloperidol, which both diminish or block the effects of psychedelics[24–26], have been shown to increase the strength of gap junction coupling[27]. Finally, psychedelics are known to bear a number of striking resemblances to the antimalarial drug mefloquine, which, like psychedelics, is a potent agonist of the 5-HT2a receptor[22], reduces low-frequency electroencephalography power[28], and can induce hallucinations and other psychiatric events[29]; importantly, mefloquine is also known to block several connexins[30–32] and is used to block gap junction coupling in experimental settings[33]. Thus, based on both our mean-field modeling results and prior empirical literature, we predict that psychedelics moderately block cortical gap junction coupling - a prediction that will need to be tested in future work.
